# Bispecific antibodies with broad neutralization potency against SARS-CoV-2 variants of concern

**DOI:** 10.1101/2024.05.05.592584

**Authors:** Adonis A. Rubio, Viren A. Baharani, Bernadeta Dadonaite, Megan Parada, Morgan E. Abernathy, Zijun Wang, Yu E. Lee, Michael R. Eso, Jennie Phung, Israel Ramos, Teresia Chen, Gina El Nesr, Jesse D. Bloom, Paul D. Bieniasz, Michel C. Nussenzweig, Christopher O. Barnes

## Abstract

The ongoing emergence of SARS-CoV-2 variants of concern (VOCs) that reduce the effectiveness of antibody therapeutics necessitates development of next-generation antibody modalities that are resilient to viral evolution. Here, we characterized N-terminal domain (NTD) and receptor binding domain (RBD)-specific monoclonal antibodies previously isolated from COVID-19 convalescent donors for their activity against emergent SARS-CoV-2 VOCs. Among these, the NTD-specific antibody C1596 displayed the greatest breadth of binding to VOCs, with cryo-EM structural analysis revealing recognition of a distinct NTD epitope outside of the site i antigenic supersite. Given C1596’s favorable binding profile, we designed a series of bispecific antibodies (bsAbs) termed CoV2-biRNs, that featured both NTD and RBD specificities. Notably, two of the C1596-inclusive bsAbs, CoV2-biRN5 and CoV2-biRN7, retained potent *in vitro* neutralization activity against all Omicron variants tested, including XBB.1.5, EG.5.1, and BA.2.86, contrasting the diminished potency of parental antibodies delivered as monotherapies or as a cocktail. Furthermore, prophylactic delivery of CoV2-biRN5 significantly reduced the viral load within the lungs of K18-hACE2 mice following challenge with SARS-CoV-2 XBB.1.5. In conclusion, our NTD-RBD bsAbs offer promising potential for the design of resilient, next-generation antibody therapeutics against SARS-CoV-2 VOCs.

**One Sentence Summary:** Bispecific antibodies with a highly cross-reactive NTD antibody demonstrate resilience to SARS-CoV-2 variants of concern.

## Introduction

Since the emergence of severe acute respiratory coronavirus 2 (SARS-CoV-2) in 2019, the COVID-19 pandemic has affected over a billion individuals globally, leading to the loss of at least 6.8 million lives (*1*). While authorized vaccines against COVID-19 have proven to substantially mitigate severe disease, there is a pressing need for additional treatment options, especially for vulnerable groups such as immunocompromised individuals. During the earlier phases of the pandemic, monoclonal antibodies (mAbs) were proven to be safe and effective as prophylactic and therapeutic modalities (*2–9*), offering significant advantages over traditional antiviral medications, including a longer half-life and a reduced incidence of side effects. However, the phenomenon of antigenic drift, especially in RNA viruses, can reduce the effectiveness of these antibody therapies over time (*10–13*). Thus, there is a critical need for next-generation antibody therapeutics capable of maintaining broad neutralizing potency despite ongoing viral evolution.

The SARS-CoV-2 spike (S), a trimeric glycoprotein on the viral surface, comprises S1 and S2 subunits responsible for binding to target cells and facilitating viral-host membrane fusion, respectively (*14–18*). Numerous neutralizing antibodies (nAbs) targeting the receptor binding domain (RBD), N-terminal domain (NTD), and subdomain 1 (SD1) in the S1 subunit have been isolated from convalescent COVID-19 donors and extensively characterized (*19–43*). In particular, RBD-specific nAbs have been shown to be the most potent and have exclusively been deployed as approved therapeutics (*44, 45*). However, with the emergence of the highly divergent Omicron variants of concern (VOCs) in 2021, the efficacy of most RBD-specific nAb clinical treatments diminished (*46–52*), leading to their withdrawal from the market. As of April 2024, only one antibody therapeutic, pemivibart, has emergency use authorization from the Food and Drug Administration for use as pre-exposure prophylaxis, representing a limited repository of approved antibody therapeutics for the treatment of SARS-CoV-2 compared to first-generation antibody therapeutics.

To address the diminished effectiveness of nAb monotherapies and therapeutic cocktails, several research groups, including ours, have developed multivalent constructs such as bispecific antibodies (bsAbs). These bsAbs feature two different antigen-binding fragments (Fabs) targeting non-overlapping spike epitopes (*41, 53–68*) and exhibit increased resilience against viral evasion compared to monoclonal antibodies, akin to antibody cocktails (*69, 70*). In addition, bsAbs offer a distinct advantage by being manufactured for delivery as a single molecule and have the potential to leverage potential avidity-driven intra-spike and inter-spike crosslinking (*54, 68*). While previous studies have explored bsAbs with Fab arms specific for the RBD and NTD (*41*), the inclusion of NTD nAbs that primarily bind to an epitope susceptible to mutations in VOCs (known as the site i antigenic supersite) (*39, 40, 71, 72*), raises concerns about potential immune evasion in current NTD-RBD bsAbs.

We recently identified NTD-specific nAbs directed to epitopes outside the site i antigenic supersite that exhibit potent neutralization and broad binding specificity against early VOCs (*20*). Given the low immune pressure on epitopes outside the antigenic supersite on the NTD, we hypothesized that a non-NTD supersite nAb could prove effective against the latest Omicron VOCs when integrated into an NTD-RBD bsAb.

Here, we conducted a comprehensive analysis of non-NTD supersite nAbs alongside an NTD supersite nAb, C1533, to assess their potency and breadth against recent Omicron VOCs. One non-NTD supersite nAb, C1596, exhibited broad cross-reactivity in binding across NTD proteins, including those from XBB.1.5, EG.5.1, and BA.2.86 VOCs. To gain insights into the breadth of C1596, we determined a 3.0 Å structure of the C1596 Fab bound to the SARS-CoV-2 S 6P trimeric glycoprotein using cryogenic electron microscopy (cryo-EM). Utilizing this structural information, we engineered SARS-CoV-2 bispecific antibodies to the RBD and NTD, referred to as CoV2-biRNs, to evaluate the efficacy of bsAbs comprising C1596 and an RBD arm. Remarkably, we identified C1596 bsAbs capable of restoring the *in vitro* neutralization effectiveness of C952 (*73*), a robust RBD nAb that had lost its efficacy against recent Omicron VOCs. Two bsAbs, CoV2-biRN5 and CoV2-biRN7, demonstrated potent neutralization against XBB.1.5, HV.1, and BA.2.86 in an *in vitro* pseudovirus neutralization assay. Additionally, when delivered prophylactically, CoV2-biRN5 significantly reduced the viral load within the lungs of K18-hACE2 mice (*74, 75*) following a challenge with SARS-CoV-2 XBB.1.5. Collectively, our data support the potential use of NTD-RBD bsAbs as a therapeutic strategy against emerging SARS-CoV-2 VOCs.

## Results

### Subhead 1: NTD-specific antibody C1596 exhibits broad cross-reactivity to SARS-CoV-2 variants of concern

Previously, we determined structures of NTD nAbs C1520, C1717, and C1791, representing antibody classes that bind epitopes outside of the site i antigenic supersite (*20*). To further compare the binding of these nAbs to a representative antibody that binds the NTD antigenic supersite, we determined a structure of SARS-CoV-2 S 6P (*76*) bound to the fragment antigen-binding (Fab) region of C1533 using single-particle cryo-EM (Figure 1A, Supplemental Figure 1A-D, and Supplemental Table 3). Similar to previously described supersite nAbs (*39, 71*), C1533 adopts a binding pose that allows engagement with the supersite β-hairpin (NTD residues 140-158) and supersite loop (NTD residues 245-264) (Supplemental Figure 1E-F). Owing to the immune pressure previously associated with the NTD antigenic supersite (*39, 40, 71, 72*), cross-reactive binding of C1533 to VOCs was limited to Wuhan-Hu-1 and Gamma NTD proteins as evaluated by enzyme-linked immunosorbent assays (ELISAs) (Figure 1B). By comparison, NTD-specific antibodies shown to recognize epitopes outside the antigenic supersite broadly recognize SARS-CoV-2 VOCs and zoonotic RaTG13 NTD proteins (Figure 1B, Supplemental Figure 2).

**Fig. 1.**
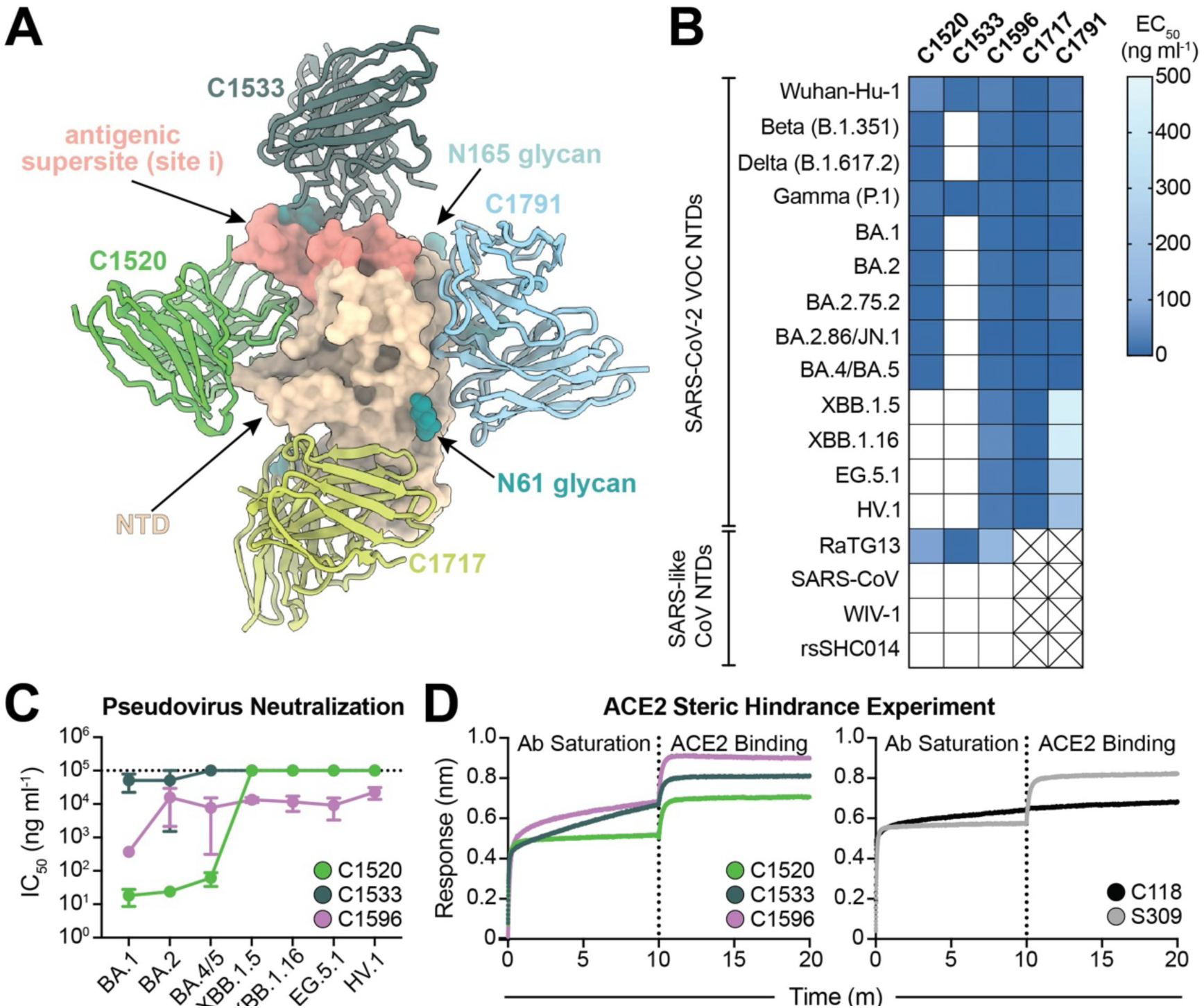
C1596, an NTD-specific mAb, exhibits broad binding and neutralizing potential across SARS-CoV-2 Omicron VOCs. **(A)**. Superimposition of C1520 (PDB: 7UAQ, green), C1717 (PDB: 7UAR, lime-yellow), C1791 (blue), and C1533 (dark slate gray) V_H_ and V_L_ domains onto the SARS-CoV-2 Wuhan-Hu-1 NTD (wheat) for a composite figure. **(B)** Heatmap of ELISA EC_50_ values of NTD-specific monoclonal IgG antibodies binding to recombinant NTD protein of SARS-CoV-2 VOCs (top) and SARS-like CoVs (bottom). Boxes shaded white represent EC_50_ values above 500 ng ml^-1^; boxes with an X indicate proteins that were not tested. Data points are represented by the geometric mean of at least three independent experiments. **(C)** Graph with IC_50_ values of NTD-specific IgG antibodies against the indicated SARS-CoV-2 VOCs. Reported data are represented by the mean of at least three replicate experiments and the standard error of the mean. **(D)** BLI association curves for soluble human ACE2 binding to SARS-CoV-2 Wuhan-Hu-1 S 6P trimer after saturation with (left) NTD IgGs and (right) RBD IgGs. S309 IgG, an RBD-specific antibody (*43*) that does not compete with ACE2 for binding (*43*), is used as a control for successful ACE2 binding following antibody saturation of the trimer; C118 IgG, an RBD-specific antibody (*23*) that competes with ACE2 for RBD binding (*42*), is used as a control for competition with ACE2.

Remarkably, C1596, an antibody that binds an epitope overlapping with that of C1791 based on epitope binning experiments (*20*), bound all tested SARS-CoV-2 VOC NTD proteins, including XBB.1.5, HV.1, and BA.2.86 with 50% effective concentration (EC_50_) values below 100 ng ml^-1^. While C1717, C1791, and C1596 exhibited a similar breadth of binding, C1596 displayed enhanced binding to the more divergent Omicron VOCs relative to C1791 and was shown to have increased neutralization potencies against pre-Omicron VOCs relative to C1717 (*20*). Consequently, we chose to further assess C1596 binding properties, comparing it with C1533 and C1520, antibodies that exhibited reduced binding breadth against pre-Omicron and late Omicron VOCs, respectively, relative to the Wuhan-Hu-1 NTD construct (Supplemental Table 1). Biolayer interferometry (BLI) experiments confirmed observations by ELISAs, showing that C1596 binds all NTDs with nanomolar affinities (Supplemental Table 1).

We next assessed the neutralization potencies of C1520, C1533, and C1596 using a SARS-CoV-2 pseudovirus-based neutralization assay (*77*) in HeLa-hACE2-TMPRSS2 cells (Figure 1C). The 50% inhibitory concentration (IC_50_) values for these mAbs against SARS-CoV-2 Wuhan-Hu-1 and early VOC pseudoviruses have been reported previously (*20*); therefore, we evaluated neutralization against recent Omicron variants. Consistent with the binding data, C1596 neutralized all pseudoviruses tested, albeit with varying strength (IC_50_ values between 0.08 and 18 µg ml^-1^), contrasting C1533 and C1520 mAbs that showed weak or no neutralizing activity against newer Omicron variants (Figure 1C, Supplemental Table 4). Finally, to probe the mechanism by which these NTD-specific antibodies mediate viral neutralization, we evaluated the ability of these mAbs to block ACE2 binding to RBD using BLI (Figure 1D). None of the tested mAbs blocked binding of immobilized SARS-CoV-2 S 6P to soluble ACE2 (Figure 1D, left panel), showing a similar profile as S309 IgG (*43*), an RBD-specific mAb that recognizes an epitope outside of the ACE2 binding motif (*19*) (Figure 1D, right panel), suggesting that these NTD Abs neutralize via a mechanism independent of blocking ACE2 engagement with the RBD. Collectively, these findings demonstrate that C1596 is highly cross-reactive against the newest Omicron VOCs, offering a potential role for non-antigenic supersite NTD mAbs in neutralizing Omicron lineages.

### Subhead 2: C1596 recognizes a quaternary epitope on the SARS-CoV-2 Spike protein comprising of the NTD, RBD, and SD1

To investigate the structural basis of C1596’s broad cross-reactivity to SARS-CoV-2 VOCs, we determined the 3.0 Å resolution structure of SARS-CoV-2 S 6P (*76*) bound to C1596 Fab using single-particle cryo-EM (Figure 2, Supplemental Figure 3, and Supplemental Table 3). The global C1596-S 6P map revealed the presence of a homogenous ternary complex with three C1596 Fabs bound to the NTDs of the S glycoprotein, which was observed in a single conformation with all RBDs in the “up” state (Figure 2A). Our structural model demonstrated that C1596 (*IGHV1-69*04/IGKV3-20*01*) primarily binds a glycopeptidic NTD epitope located between the N61_NTD_ and N234_NTD_ glycans (∼64% of the total epitope buried surface area (BSA)), while also engaging the RBD and subdomain 1 (SD1) of the same protomer, which contribute 23% and 14% to the total epitope BSA, respectively (Figure 2B-C, Supplemental Figure 4A, and Supplemental Table 2).

**Fig. 2.**
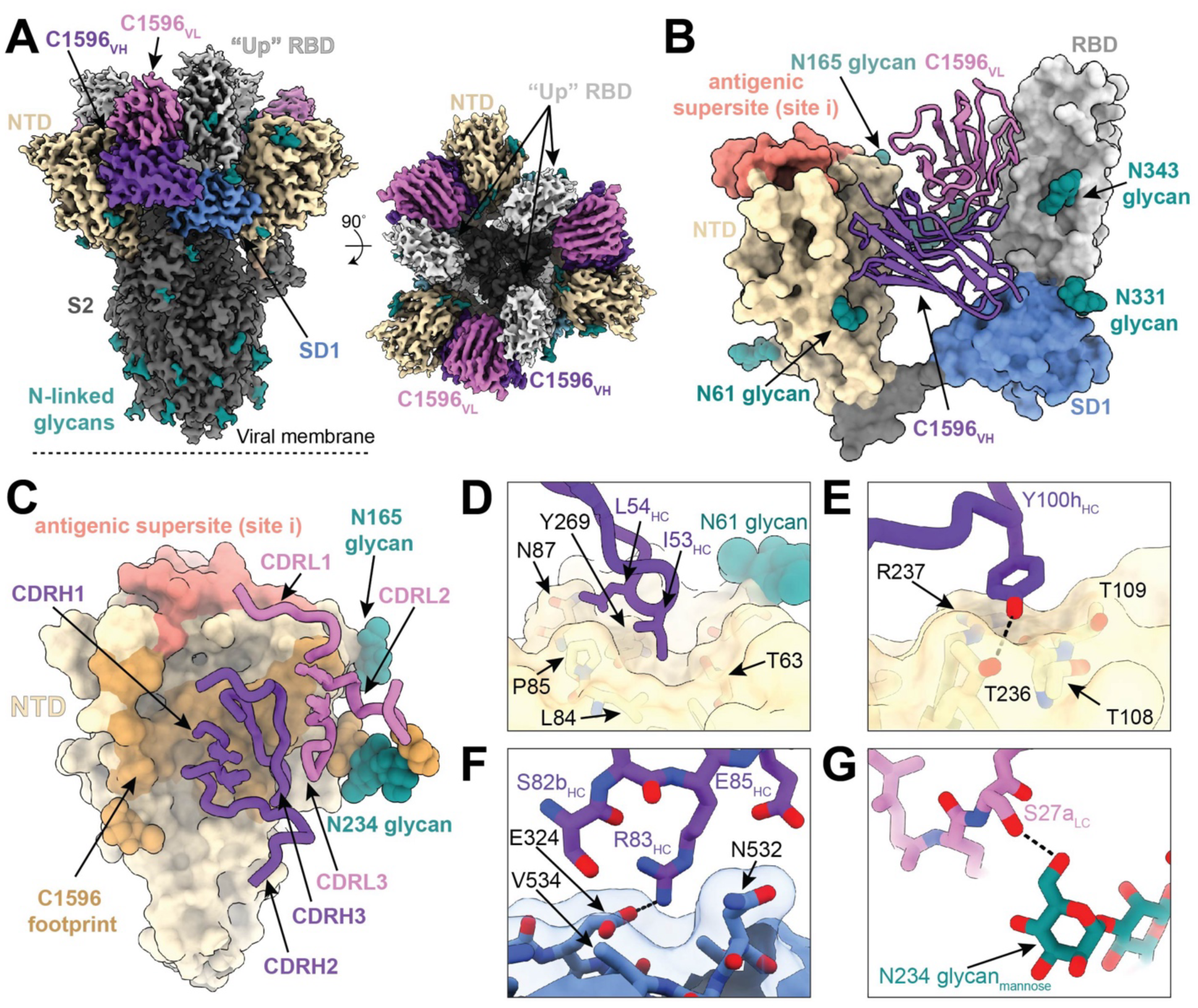
C1596 binds the N-terminal domain of the SARS-CoV-2 Spike protein outside of the antigenic site i supersite. **(A)** 3.0 Å cryo-EM density for C1596-S 6P trimer complex; C1596 variable domains (shades of purple) bound to the NTD (wheat), with neighboring RBD (light gray) and SD1 (cornflower blue). **(B)** Structural model of C1596 (shades of purple, ribbon) binding to the NTD (wheat), in an epitope distinct from the antigenic supersite (salmon), with additional contacts with the RBD and SD1. **(C)** C1596 footprint (brown), as defined by buried surface area, depicted on a surface representation of the NTD (wheat) with CDR loops of the C1596 heavy (purple) and light chains (pink) annotated. **(D)** C1596 CDRH2 residues I53-L54 target a hydrophobic pocket at the base of the N61-glycan on the NTD. **(E)** C1596 CDRH3 residue Y100h engagement with a polar surface on the NTD. **(F)** Potential salt bridge mediated by the C1596 residue R83 and NTD residue E324 in the SD1 (cornflower blue). **(G)** Representation of a potential hydrogen bond between the C1596 LC (pink) and the N234 glycan mannose (teal) of the NTD. Hydrogen bonds were annotated as defined by PDBePISA (see Methods).

To investigate the importance of the contacts in the RBD and SD1, we evaluated the binding profiles of C1596 to SARS-CoV-2 S 6P compared to recombinant NTD and RBD-SD1 proteins (Supplemental Figure 4B-F). While the overall binding affinities to Wuhan-Hu-1 S 6P and NTD were similar (100 nM and 160 nM, respectively), the dissociation rate of C1596 from the NTD was ∼10-fold faster than from S 6P (Supplemental Figure 4B-C). Given the slower dissociation kinetics observed for binding to S 6P, we assessed the binding of C1596 to the RBD or RBD-SD1. Despite approximately one-third of the epitope involving interactions with the RBD and SD1 domains, BLI analyses showed no appreciable binding, indicating that contacts in the RBD and SD1 alone are not sufficient to sustain binding to C1596 (Supplemental Figure 4D-E). Additionally, the binding affinity of C1596 to SARS-CoV-2 BA.1 S 6P, which consists of mutations S371, S373P, and S375F in the RBD that sit adjacent to interfacing residues (Y369, A372, and F377; Supplemental Figure 4A), was similar to Wuhan-Hu-1 S 6P, suggesting a limited role for RBD mutations in mediating viral escape (Supplemental Figure 4B and 4F). However, RBD “up” states are necessary for C1596 binding to prevent steric clashes with the RBD on adjacent spike protomers, as illustrated by the inability of C5196 to bind to a “closed” spike trimer via BLI (Supplemental Figure 4G-I).

Analysis of the Fab-NTD interface revealed that the C1596 heavy chain (HC) primarily mediates NTD contacts (C1596 HC represented ∼912 Å^2^ of ∼1264 Å^2^ paratope BSA), which comprises residues along the N1 loop (residues 22-28), the β3 strand (residues 63 and 65), the β6 strand (residues 82-88), the hairpin turn between the β8 and β9 strands (residues 108 and 109), the β17 strand (residues 235-237), and the β19 strand (residues 267 and 269) (Figure 2C, Supplemental Table 2). Interestingly, the C1596 CDRH2 loop is buried into a hydrophobic pocket along the surface of the NTD using residues I53_HC_ and L54_HC_ (Figure 2D), a germline-encoded motif in IGHV1-69 broadly neutralizing antibodies previously identified as an anchor for interactions to conserved hydrophobic epitopes in influenza, HIV-1, and hepatitis C viral antigens (*78*). CDRH3 residues 97 to 100_B_ account for ∼21% of the total HC paratope BSA, packing into a groove along the NTD comprising the β6 and β17 strands, while residue Y100_H HC_ engages a polar surface comprising NTD residues T108_NTD_, T109_NTD_, T236_NTD_, and R237_NTD_ (Figure 2E and Supplemental Figure 4J). Additional contacts by the CDRH1 loop and framework region (FWR) 3, including a potential salt bridge between R83_C1596 HC_ and E324_SD1_, account for the remainder of HC interactions (Figure 2F and Supplemental Figure 4K). The C1596 light chain accounts for ∼28% of the total BSA of the paratope and primarily engages the RBD and the N234_NTD_ glycan, which has been previously shown to serve as a linchpin for RBD “up” states (*79*) (Figure 2C,G and Supplemental Table 2).

Given C1596’s broad footprint that spans three domains within the same S1 protomer, we next assessed the overall contribution of somatic hypermutations (SHMs) in the V gene-encoded region of the antibody paratope (Supplemental Figure 5A-B). While the V_H_ domain had six SHMs, only two of eighteen total residues comprising the heavy chain paratope involved SHMs, whereas all V_L_ domain paratope residues were germline encoded, suggesting that similar to other nAbs isolated following infection (*23*), C1596 has minimal somatic mutations involved at the contacting interface. Finally, we compared the binding pose of C1596 to C1791 (*IGHV3-23*01/IGKV1-17*01*), an antibody described by our group (*20*), and S2L20 (*IGHV3-30/IGKV1-33*) (*39*), a previously-described cross-reactive, but weakly neutralizing (IC_50_ > 1 ug ml^-1^) antibody (*80*) (Supplemental Figure 5C-E). While the NTD footprint was similar across all antibodies and each interacted with the SD1 domain, available structures showed that only C1596 engaged the RBD within the same protomer (Supplemental Figure 5E). Thus, despite 15 shared NTD contacts between mAbs in this class, stabilizing interactions against the “up” RBD by C1596 likely explain its increased binding against later VOCs such as Omicron XBB.1.5, relative to C1791 (Figure 1B). Overall, our structural analysis demonstrates that C1596 predominantly interacts with a previously-described NTD glycopeptidic epitope (*20, 39*), with additional stabilizing contacts in the RBD and SD1, making it distinguishable from other antibodies in its class.

### Subhead 3: C952-C1596 bispecific antibodies neutralize SARS-CoV-2 pseudotyped variants of concern

Previous studies have highlighted the effectiveness of bispecific constructs (*41, 53–68*), including one notable example where constructs paired broadly cross-reactive weak or non-neutralizing mAbs with a potent inhibitor of viral entry, which significantly enhanced its overall neutralization potency and breadth (*55*). Given C1596’s broad cross-reactivity against SARS-CoV-2 VOCs and trapping of RBDs in an “up” state, we investigated the ability of C1596 to support bivalent interactions with an RBD-specific antibody in a single-molecule bispecific format (Supplemental Figure 6). Utilizing a cut-off distance of ∼65 Å between the Cα atoms of residues near the C-termini of adjacent Fab C_H_1 domains (*19*), we speculated that C1596 could likely pair with a class 3 RBD-specific mAb (*19*) to support bivalent binding within a single Spike (Supplemental Figure 6C). We previously identified a subset of broad and potent class 3 RBD-specific mAbs from donors one-year post-SARS-CoV-2 infection (*73*). Of these antibodies, C952 was exceptionally potent and resistant to mutations identified in early VOCs.

To investigate the basis of potent C952-mediated inhibition of viral entry and broad binding specificity to VOCs (Figure 3A), we determined a single-particle cryo-EM reconstruction at 3.4 Å for a C952-S 6P complex (Figure 3B and Supplemental Figure 7). As also observed in a previous structure of the clonal ancestor C032 (*81*), Fabs of C952 bound both “up” and “down” RBD states, recognizing a conserved glycopeptidic RBD epitope that includes residues 439-451 along the RBD ridge and residues at the base of the N343_RBD_ glycan (Figure 3C,D). Consistent with the C032 structure, C952 CDRL3 and CDRH3 loops recognize residue R346_RBD_ and a loop comprising residues 440-450, respectively, which are RBD regions commonly mutated among VOCs (*82*) (Figure 3E). Yet, increased SHM in the C952 paratope, which was also observed for the 6-month clonal ancestor C080 (*81*), likely explains resistance to single-point mutations (*73*) and Omicron VOCs that harbor multiple mutations in these regions (Figure 3A). Interestingly, unlike canonical class 3 mAbs (*19*), C952 adopts a pose where CDRH1 loop residues are modeled to potentially clash with ACE2-bound RBD (Supplemental Figure 7E). Indeed, BLI competition analyses revealed diminished binding of soluble ACE2 to immobilized Spike trimers in the presence of C952, which was not observed for class 3 mAbs C135 and S309 (Supplemental Figure 7F), likely explaining potent C952-mediated neutralization.

**Fig. 3.**
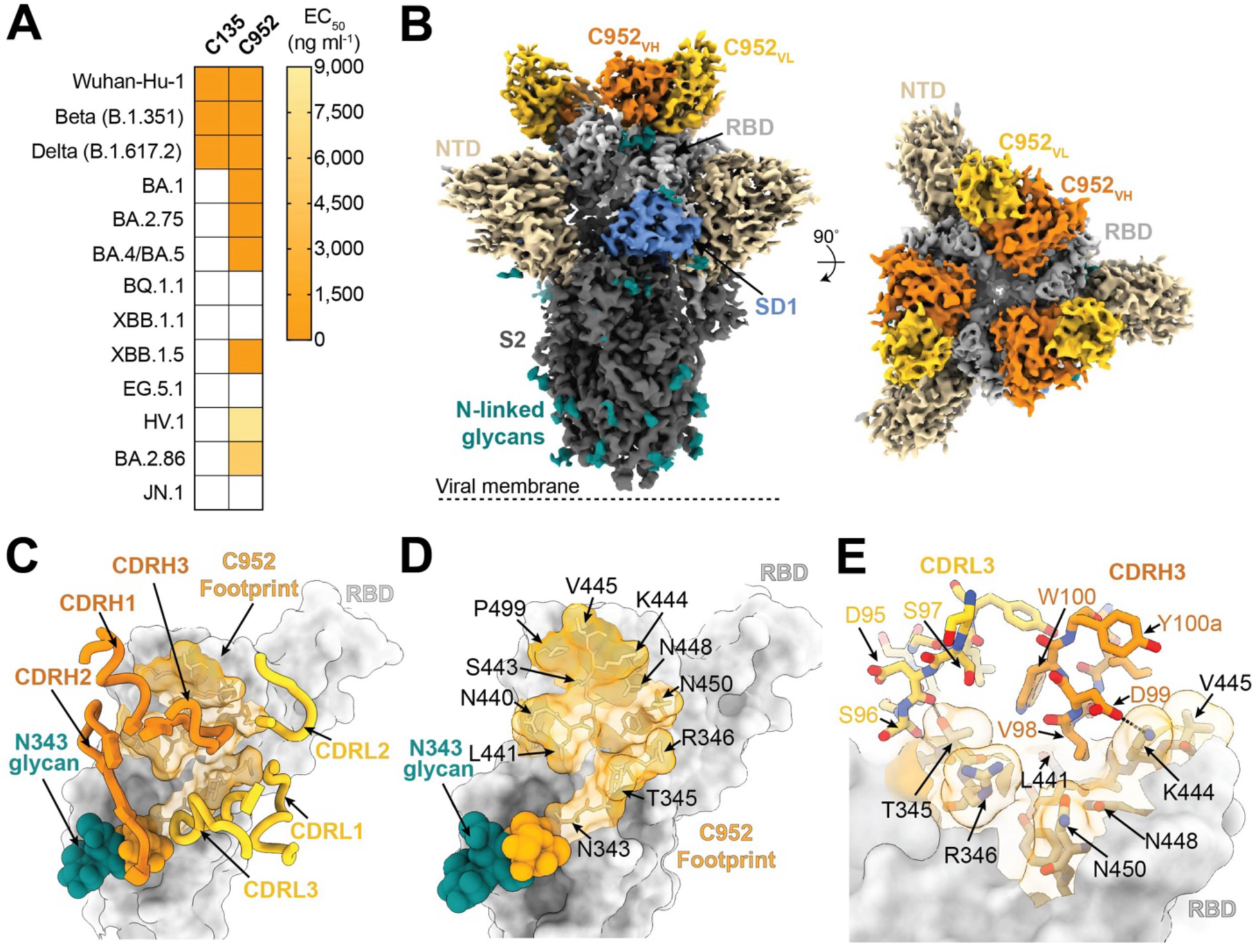
C952 binds a class 3 epitope of the SARS-CoV-2 Spike RBD. **(A)** Heatmap of ELISA EC_50_ values of C135 and C952, two previously characterized RBD-specific monoclonal IgG antibodies (*23, 73*), binding to recombinant RBDs of SARS-CoV-2 VOCs. Boxes shaded white represent EC_50_ values above 9,000 ng ml^-1^. Data points are represented by the geometric mean of at least three independent experiments. **(B)** 3.4 Å cryo-EM density for C952-S 6P trimer complex; C952 V_H_ (orange) and V_L_ (yellow) bound to the RBD (light gray), with neighboring NTD (wheat) and SD1 (cornflower blue). **(C-D)** C952 footprint (light orange), as defined by buried surface area, depicted on a surface representation of the RBD (light gray) with **(C)** CDR loops of the C952 V_H_ (dark orange) and V_L_ (yellow) and **(D)** RBD epitope residues denoted. **(E)** CDRH3- and CDRL3-mediated contacts on the RBD. Potential hydrogen bonds are depicted by black dashed lines.

To confirm whether the binding modes and distances of C952 and C1596 would be compatible in an IgG-like bispecific that promotes bivalent interactions within a single Spike, we modeled Fab binding to the same protomer of the trimeric SARS-CoV-2 Wuhan-Hu-1 S protein (Figure 4A). Modeling revealed Cα distances between C_H_1 C-terminal residues of the two Fabs of ∼34 Å, indicating their potential pairing in an IgG-like bispecific. However, minor steric clashing between the C1596 V_L_ and C952 V_H_ FWRs in our model was observed (Figure 4A). Thus, we performed a BLI competition assay to assess the possibility of both Fab specificities binding to the same Spike trimer. In this assay, immobilized S 6P was first bound by C952 or C1596 Fab, and the complex was subsequently assayed for binding to the Fab belonging to the other specificity (Figure 4B). Our data indicated no steric hindrance when either Fab was associated with the S trimer first, validating their incorporation in a bispecific format.

**Fig. 4.**
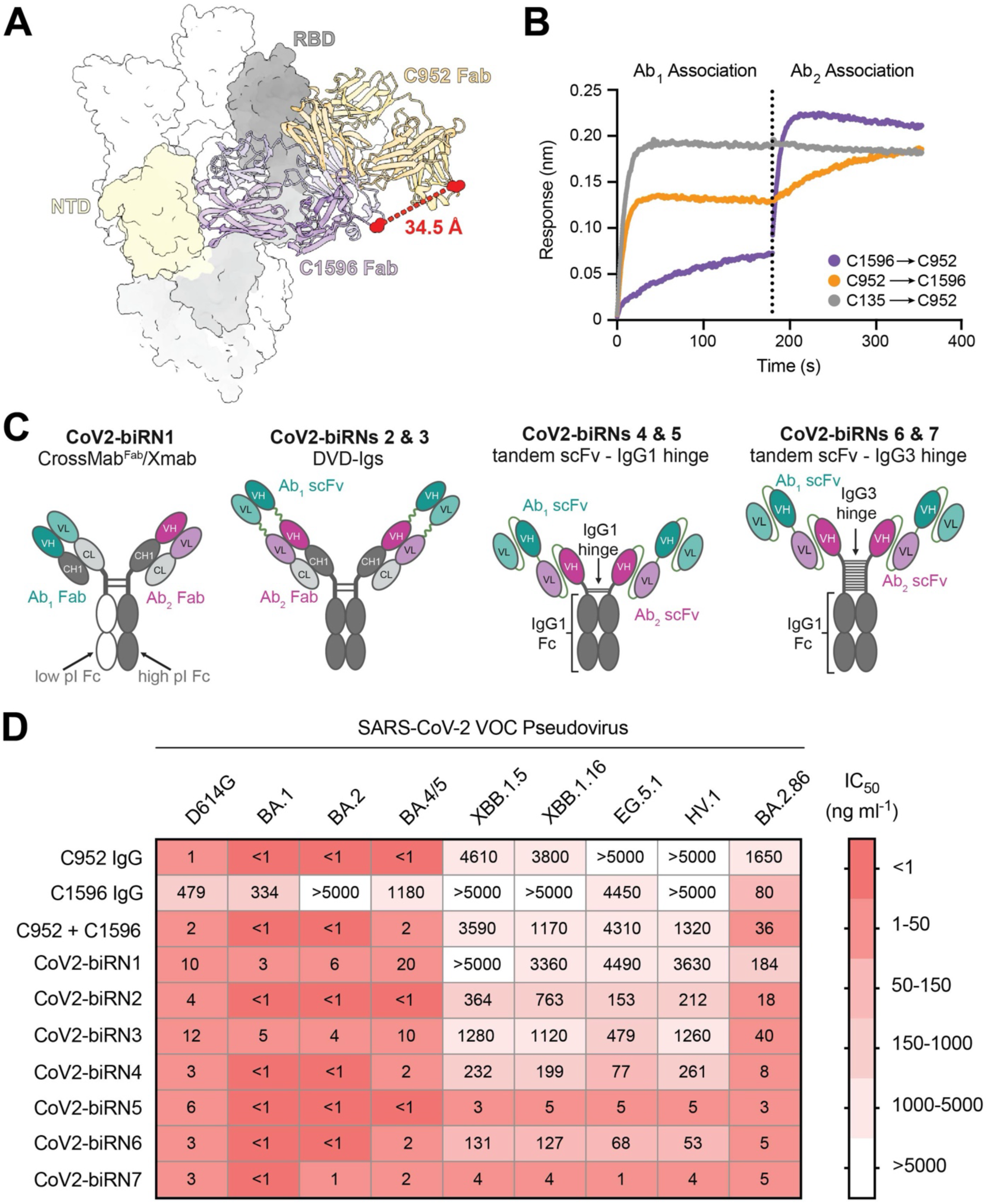
CoV2-biRN5 and CoV2-biRN7 exhibit robust *in vitro* neutralization efficacy against Omicron VOCs. **(A)** Structural modeling of C1596 and C952 Fabs bound to SARS-CoV-2 S 6P. Distance between the C-terminal residue of the C_H_1 domain is depicted by the red dotted line. **(B)** BLI experiment testing simultaneous binding of either C952 and C1596 Fabs to SARS-CoV-2 S 6P trimer (Ab_2_ association) when the S trimer is saturated with the other Fab (Ab_1_ association). C135, an anti-RBD antibody (*23*), is depicted as a control to demonstrate steric clashing with C952. **(C)** Schematic of CoV2-biRN constructs. (See Supplemental Figure 8A for details of constructs) **(D)** Heatmap of IC_50_ values of parental C952 and C1596 IgGs, a cocktail of C952 and C1596 IgGs, and CoV2-biRNs against SARS-CoV-2 VOC pseudoviruses. Reported values are represented by the geometric mean of at least three replicate experiments.

Thus, we next designed several IgG-like bispecific constructs, termed CoV2-biRN1-7, that utilized C1596 and C952 specificities (Figure 4C and Supplemental Figure 8A). These constructs included CoV2-biRN1, a traditional IgG-like bispecific based on CrossMab and Xmab bispecific platforms (*83, 84*); CoV2-biRNs 2 and 3, bsAb constructs utilizing the dual variable domain immunoglobulin (DVD-Ig) format (*85*); and CoV2-biRNs 4-7, which were designed to incorporate tandem single-chain fragment variable domains (scFvs) with either an IgG1-hinge (CoV2-biRNs 4 and 5) or IgG3-hinge (CoV2-biRNs 6 and 7), the latter of which designed to afford increased flexibility of the bsAb arms observed for IgG3 antibodies (*86*). Following expression and purification of the bsAb constructs (Supplemental Figure 8B-D), we next compared the *in vitro* neutralization potencies of CoV2-biRNs 1-7 relative to the parental nAbs alone and as a cocktail (Figure 4D and Supplemental Table 4). The IC_50_ values of CoV2-biRN1 were comparable to or weaker than the cocktail of the parental antibodies, whereas CoV2-biRNs 2 and 3 demonstrated minor improvements in neutralization potency with IC_50_ values ranging from 150-1,300 ng ml^-1^ against the latest Omicron VOCs (Figure 4D). CoV2-biRNs 4-7 (tandem scFv constructs) demonstrated enhanced neutralization activity in this assay relative to the parental cocktail, with IC_50_ values less than 300 ng ml^-1^ across all VOC pseudoviruses tested. Notably, CoV2-biRNs 5 and 7, tandem scFv bsAbs with the C1596 scFv as the outer domain and either the IgG1 or IgG3 hinge, respectively, retained potency (IC_50_ values < 10 ng ml^-1^) across all Omicron VOCs, including the divergent BA.2.86 strain (Figure 4D). To assess whether a bispecific with an NTD supersite nAb (C1533) paired with C952 would have similar efficacy, we engineered CoV2-biRN8 and CoV2-biRN9 and evaluated their neutralization potencies (Supplemental Figure 8E,F). As expected, CoV2-biRN5 demonstrated stronger neutralization compared to both CoV2-biRN8 and CoV2-biRN9, highlighting the importance of tethering bsAbs using an NTD arm with breadth of binding and reinforcing the advantage of utilizing a non-supersite antibody. Collectively, our results illustrate the ability to engineer two bsAbs, CoV2-biRN5 and CoV2-biRN7, that retain potent neutralization activity against the latest Omicron VOCs.

### Subhead 4: Full-length spike pseudovirus deep mutational scanning reveals infrequent escape mutations in the NTD in response to CoV2-biRN5

To understand how spike mutations affect neutralization by CoV2-biRN5, we utilized a full-length SARS-CoV-2 XBB.1.5 pseudovirus-based deep mutational scanning library to map escape mutations as described (*87, 88*) (Figure 5). The escape mutational profile for CoV2-biRN5, characterized by the log_2_ fold change in neutralization IC_90_ values, mapped to both the NTD (β6 and β17 strands) and RBD (ridge loop residues 440-450), contrasting the mutational escape profile against the parental cocktail that focused mainly on residues with the C952 RBD epitope (Figure 5A,B). These findings underscore the critical role of the NTD antibody arm in inhibiting viral entry in the bispecific construct, particularly considering that the parental cocktail has a neutralization profile similar to that of C952 alone, which likely drives the neutralization potency in the parental cocktail. Among the escape mutations identified in the NTD for CoV2-biRN5, three showed greater than 50-fold change in IC_90_ values: P85del, N87I, and R237Y; residues that comprise an NTD groove, which is engaged by the C1596 CDRH3 (Figure 2E).

**Fig. 5.**
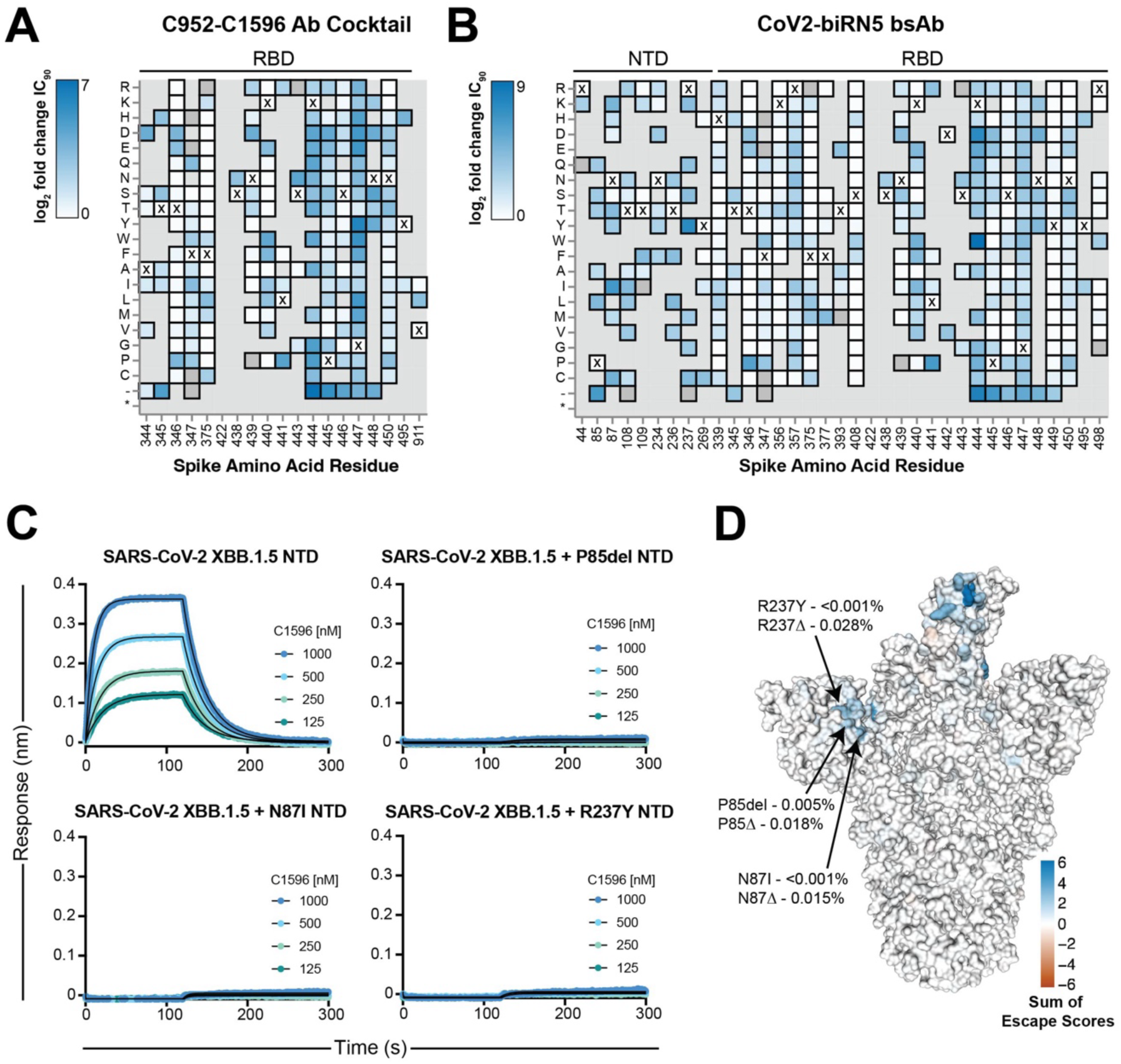
Deep mutational scanning of SARS-CoV-2 XBB.1.5 Spike and C952-C1596 antibody cocktail and CoV2-biRN5 escape. **(A)-(B)** Heatmap of log_2_ fold change in IC_90_ of greater than 3.4 for residues in the full-length SARS-CoV-2 XBB.1.5 Spike in a pseudovirus assay with **(A)** an equimolar cocktail of C952 and C1596 IgGs or **(B)** CoV2-biRN5. Residues marked with an X represent the amino acid residue present in the XBB.1.5 VOC Spike; amino acid mutations that are absent from the library are depicted in gray. Complete analysis and code for deep mutation scanning can be found at https://github.com/dms-vep/SARS-CoV-2_XBB.1.5_spike_DMS_Barnes_mAbs. **(C)** BLI association and dissociation curves, illustrating the effect of escape mutations in the XBB.1.5 NTD on C1596 binding. The concentrations of C1596 tested are indicated on each panel. Dissociation phase begins at time = 120 s. Fit curves are depicted as black solid lines. **(D)** Surface rendering of SARS-CoV-2 XBB.1.5 Spike colored by the sum of escape scores at each residue. Key residues of escape labeled along with their frequency across 1,984,175 sequences uploaded to GISAID CoVsurver as of February 26, 2024; frequency is denoted for the escape mutation with the greatest escape score (top) and for all amino acid mutations at that residue site (bottom).

To validate the phenotypes of these NTD escape mutations, we introduced individual mutations into the SARS-CoV-2 XBB.1.5 NTD protein and assessed the binding of C1596 to each construct (Figure 5C). As expected, C1596 binding was completely abrogated against the three NTD variants, confirming the loss of neutralization potency observed in the deep mutational scanning library assay. Next, after identifying NTD escape mutations in this assay, we investigated the prevalence of these mutations in circulating strains of SARS-CoV-2. Using sequences deposited to the GISAID CoVsurver (*89*), we calculated the frequency of the three strongest escape mutations, as well as the frequency of any mutation, insertion, or deletion at the three escape sites (Figure 5D). Each individual mutation was found in less than 0.005% of all sequences evaluated, suggesting that the epitope targeted by C1596-like mAbs is under minimal selective pressure and that bsAbs utilizing C1596 as an anchoring modality, such as CoV2-biRN5, are likely resilient to circulating strains.

### Subhead 5: CoV2-biRN5 reduces SARS-CoV-2 XBB.15 viral load in a K18-hACE2 mouse model

Finally, to determine the *in vivo* relevance of these findings given the unusual architectures of our most potent bsAb constructs, we assessed the efficacy of CoV2-biRN5, and the parental mAb cocktail, against SARS-CoV-2 XBB.1.5 challenge using a K18-hACE2 transgenic mouse model (*74, 75*) (Figure 6A). CoV2-biRN5 or the mAb cocktail was administered intraperitoneally at a dose of 5 mg kg^-1^, either 24 hours before (prophylactic arm) or 4 hours after (treatment arm) intranasal administration of 2 x 10^4^ plaque-forming units (PFUs) of SARS-CoV-2 XBB.1.5. Given the variability in weight loss phenotypes observed in K18-ACE2 transgenic mice challenged with Omicron strains (*90*), lung tissue was harvested from the mice three days post-infection and assessed for viral load by quantifying RNA copy number. Prophylactic administration of CoV2-biRN5 resulted in a statistically significant ∼1,500-fold reduction in viral load compared to the untreated control, consistent with findings of previously EUA-approved mAbs administered at similar doses in K18-ACE2 mice (*91, 92*) (Figure 6B). While the mean viral load was ∼110-fold lower in mice treated with CoV2-biRN5 following XBB.1.5 challenge, a statistically significant fold-change was not observed at a dose of 5 mg kg^-1^ (Figure 6C). Collectively, these findings demonstrate the efficacy of CoV2-biRN5 in an animal model, thereby indicating its promising potential as a prophylaxis candidate for SARS-CoV-2 VOCs.

**Fig. 6.**
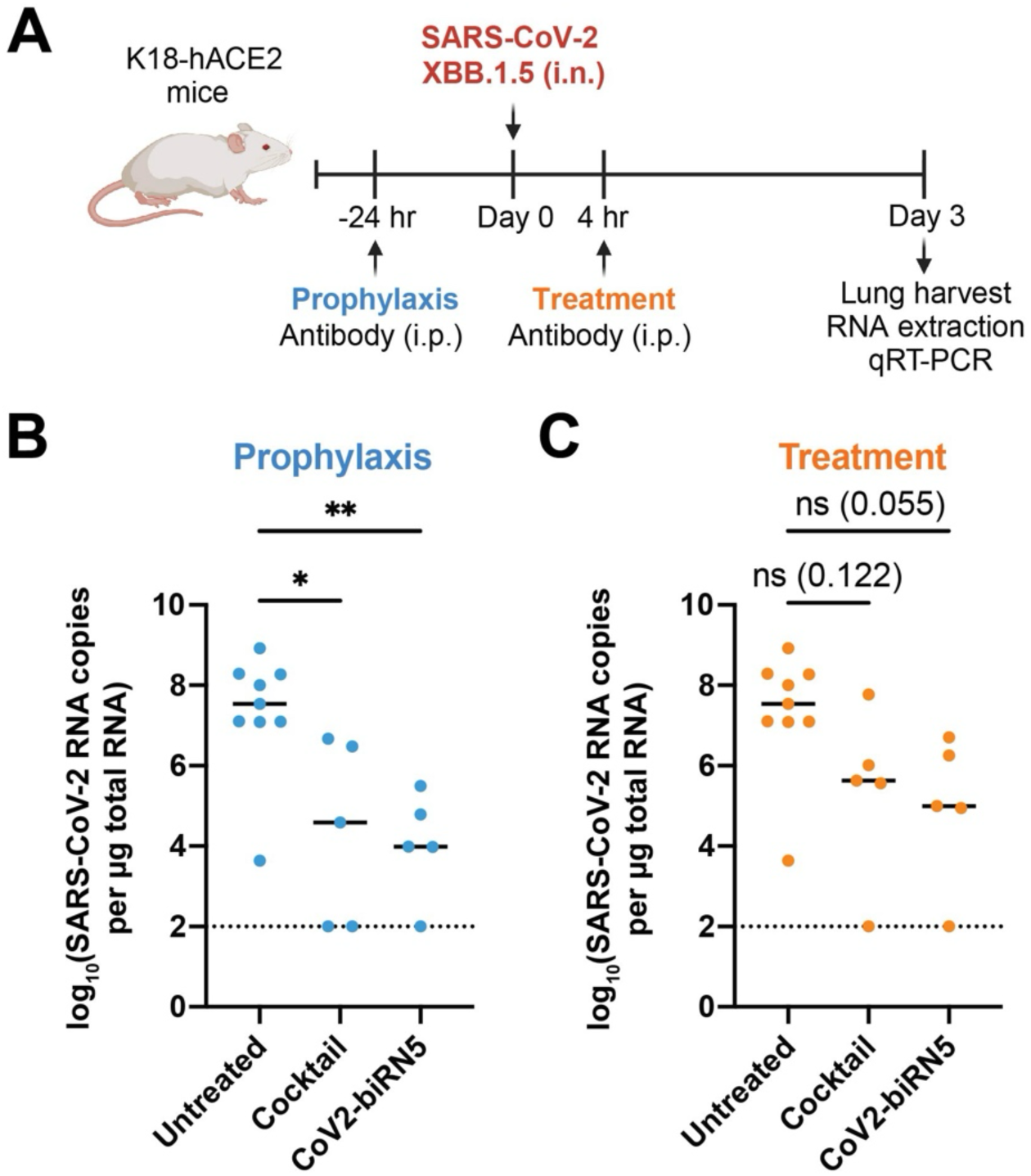
Bispecific CoV2-biRN5 demonstrates prophylactic efficacy in a K18-hACE2 mouse model. **(A)** Schematic representation of the experimental timeline for assessing the efficacy of CoV2-biRN5 *in vivo*. K18-hACE2 mice received 5 mg kg^-1^ of either the cocktail of C952 and C1596 IgGs or the CoV2-biRN5 bsAb by intraperitoneal (i.p.) injection 24 hours before or four hours after intranasal (i.n.) inoculation with 2 x 10^4^ PFU of the SARS-CoV-2 XBB.1.5 virus. Lungs were harvested at three days after infection. **(B)** and **(C)** Viral RNA load of SARS-CoV-2 XBB.1.5 in the lung was measured for each group (line indicates mean; in order from left to right, *n* = 9, 5, and 5) in the **(B)** prophylaxis and **(C)** treatment arm. Data were analyzed using one-way ANOVA, correcting for multiple comparisons with Dunnett’s test with comparison to the untreated control. NS, not significant, \*\**P*<0.01, \**P*<0.05. Dotted line indicates the lower limit of detection of the assay.

## Discussion

Protective antibody responses to SARS-CoV-2 predominantly comprise antibodies that target the RBD (*44, 45*), as expected given its role in viral entry. However, this dominant neutralizing response has contributed to immune pressure, giving rise to VOCs (*13*) that have rendered first-generation COVID-19 antibody therapeutics ineffective (*46–52*), thus limiting the therapeutic arsenal for immunocompromised individuals who are susceptible to severe disease. Thus, there remains a critical need to identify next-generation antibody therapeutics that will withstand the ongoing evolution of SARS-CoV-2 by targeting spike epitopes that are conserved among SARS-CoV-2 variants. Towards this goal, our group and others previously isolated and characterized neutralizing antibodies with appreciable breadth against VOCs that bind the spike NTD outside of the antigenic supersite (*20*). Here, we further evaluate the vast breadth of binding to recent Omicron VOCs for non-supersite NTD antibodies, including NTD nAbs C1520, C1717, C1791, and C1596.

We present the cryo-EM structure of SARS-CoV-2 S 6P bound to C1596, shedding light on its recognition of a quaternary epitope encompassing the NTD, RBD, and SD1 domains within a single spike protomer. Notably, our investigation of binding kinetics revealed that C1596 exhibits a slower dissociation rate (*k*_d_) when bound to the spike trimer compared to the monomeric NTD, which we speculate plays a critical role in C1596 resilience to spike mutations observed in recent VOCs. This resilience may stem from the inability of non-supersite NTD mAbs to engage in intraspike avidity due to the spatial orientation of epitopes on the spike trimer. By burying more Fab surface area against the RBD and SD1 within the same protomer, C1596 forms interactions which, while individually insufficient for binding, collectively contribute to its ability to maintain high affinity for the spike, even in the presence of mutations in the primary NTD epitope (e.g., XBB.1.5). This multifaceted interaction that increases C1596 affinity by slower dissociation kinetics is akin to observations observed for somatically mutated RBD-specific mAbs where mutations outside the Fab-antigen interface enhanced affinities, resulting in antibody resilience to VOCs (*81*).

In light of the broad binding spectrum and slow dissociation kinetics observed for C1596, we explored its utility in a bispecific format. Prior studies have proposed leveraging the robust binding capacity of weak or non-neutralizing antibodies within multispecific proteins, using them as critical tethers to enhance interactions with other inhibitors (*55, 64*). Hence, we reasoned that C1596 could serve as an anchoring component in bispecific antibody therapeutics, conferring dual specificity for the NTD and RBD for the prevention and treatment of SARS-CoV-2 VOCs. When coupled with the RBD-specific nAb C952, six of seven bispecific constructs exhibited heightened neutralization efficacies compared to the individual parental nAbs, either alone or in a cocktail against Omicron VOCs that emerged after BA.4/5. Interestingly, these six constructs incorporated both specificities within the same arm of an IgG-like molecule, likely resulting in increased intraspike avidity effects relative to the parental cocktail or CrossMab bsAb. Notably, among these constructs, CoV2-biRN5 and CoV2-biRN7 demonstrated sustained neutralization efficacy across all VOCs tested. These constructs were designed with C1596 positioned N-terminal to C952, differing from CoV2-biRN4 and CoV2-biRN6 (C952 at the N-terminus), which exhibited lower potencies. This observation underscores the significance of C1596’s initial association with the trimer, inducing an “up” RBD state that facilitates subsequent binding by C952. In contrast, when C952 is at the N-terminus of the construct, its capability to recognize both “up” or “down” RBD states, the latter being incompatible with C1596 binding, likely constrains benefits derived from intraspike avidity effects.

Collectively, these findings underscore the strategic importance of antibody orientation and the sequential order of binding events in designing effective bispecific therapeutics against SARS-CoV-2. As currently designed, CoV2-biRN5 has a potential prophylactic application against SARS-CoV-2. Findings from animal challenge studies, with a dose regimen of 5 mg kg^-1^, position CoV2-biRN5 alongside previously FDA-authorized monoclonal antibody treatments (*91*), underscoring its efficacy and therapeutic potential. Moreover, results from pseudovirus deep mutational scanning experiments indicate that the emergence of escape mutations for CoV2-biRN5 is currently minimal, with a frequency of less than 0.05% among available sequences of circulating SARS-CoV-2 variants. This low prevalence of naturally occurring mutations suggests that the epitope recognized by CoV2-biRN5 is not under appreciable selective pressure, further bolstering the viability of CoV2-biRN5 as an antibody therapeutic option for individuals at risk of severe disease.

While our study presents innovative findings, it is important to acknowledge several limitations to the work. First, our bsAb constructs are limited to a single antibody pairing, namely C1596 and C952, serving as a proof-of-concept for leveraging non-supersite NTD mAbs to design potent bsAbs. Future investigations could benefit from exploring additional antibody pairs, where both specificities are resilient to all VOCs, to validate and expand upon this proof-of-concept. Furthermore, although the tandem scFv bsAbs (CoV2-biRNs 4 to 7) exhibited the highest potency among the tested bsAbs, their expression and purification will need to be further optimized to increase overall yields. Addressing this issue will require additional biochemical modifications to enhance the manufacturability of these bsAbs for further investigation. Additionally, a more thorough characterization of the bioavailability, pharmacokinetics, and Fc-mediated effector functions of the bsAb compared to the parental antibodies could provide additional insights. Such insights would contribute to a better understanding of the observed similar efficacy of the bsAb and cocktail formulations in animal models, despite their divergent performance in our *in vitro* experiments. These considerations highlight avenues for future research aimed at refining and expanding the scope of our work.

In summary, our findings underscore the utility of employing a highly cross-reactive antibody as a tethering arm in a bispecific reagent, thereby restoring the potency of inhibitors that have lost neutralization efficacy due to weaker binding affinities to their epitopes. While our study focused on a specific antibody pairing, there is potential for further investigation using additional antibody combinations targeting other highly conserved epitopes on the SARS-CoV-2 S glycoprotein, such as the fusion peptide and stem helix. Moreover, the versatility of this platform extends beyond SARS-CoV-2, with potential applications against other viral proteins, including those in influenza or HIV. By leveraging broadly binding antibodies to tether other antibodies or inhibitors, these bispecific modalities offer a promising next-generation prophylactic or therapeutic option. Moving forward, preclinical studies should explore the efficacy of these bispecific antibodies in protecting against emerging SARS-CoV-2 variants of concern, paving the way for their potential clinical translation.

## Materials and Methods

### Cell lines

HEK293T cells (*Homo sapiens*, embryonic kidney cells) used for pseudovirus generation were obtained from Dr. Pamela Bjorkman (California Institute of Technology). HeLa-ACE2-TMPRSS2 cells (*Homo sapiens*, cervical epithelial cells expressing ACE2 and TMPRSS2) used for pseudovirus neutralization assays were obtained from Dr. Peter S. Kim (Stanford University). All adherent cells were cultured in D10 media (Dulbecco’s Modified Eagle Medium (DMEM) (Gibco, 11995065) supplemented with 10% heat-inactivated fetal bovine serum (FBS), 100 U/ml-100 µg/ml Penicillin-Streptomycin, and 1% 2 mM L-Glutamine) at 37°C and 5% CO_2_. Expi293F™ cells (*Homo sapiens*, embryonic kidney cells in suspension) used for transient protein expression were purchased (ThermoFisher Scientific, A14527) and cultured in Expi293™ Expression Medium (ThermoFisher Scientific, A1435104) while shaking at 37°C and 8% CO_2_.

### Monoclonal antibody cloning, expression, and purification

Sequences for S309, C118, C135, C952, C1520, C1533, and C1596 anti-SARS-CoV-2 IgG antibodies were previously published (*20, 23, 43, 73*). Heavy IgG and light chains for each antibody were cloned into the AbVec2.1 expression plasmid (gift from Dr. Pamela Bjorkman, California Institute of Technology). Fab constructs were obtained by subcloning the variable heavy (V_H_) and constant heavy chain 1 (CH_1_) domain from the IgG vectors into an AbVec2.1 expression plasmid with a C-terminal 6x histidine tag. For IgG and Fab expression, heavy and light chains were co-transfected in Expi293F cells using the ExpiFectamine transfection kit (ThermoFisher Scientific, A14525) in a 1:1 ratio; ExpiFectamine transfection enhancer was added 20 h post-transfection. IgGs and Fabs were isolated and purified from filtered cell supernatants via affinity chromatography using HiTrap MabSelect SuRe (Cytiva) and HisTrap HP (Cytiva) columns, respectively. Elutions were concentrated using Amicon™ spin concentrators and further purified by size-exclusion chromatography on a HiLoad 16/600 Superdex 200 pg (Cytiva) and Superdex 200 Increase 10/300 GL (Cytiva) columns using an AKTA pure 25 M1 system (Cytiva). Proteins were concentrated via Amicon™ spin concentrators and stored in Tris-Buffered Saline with azide (1x TBS-Az; 20 mM Tris pH 8.0, 150 mM NaCl, 0.02% Sodium Azide) at 4°C.

### Bispecific antibody cloning, expression, and purification

CoV2-biRN1 was designed and cloned utilizing the CrossMab^Fab^ and Xmab bispecific formats, as previously described (*83, 84*). Briefly, the IgG1 Fc chain was mutated to create an isoelectric point (pI) differential for the heterodimeric formation of the IgG heavy chains in the asymmetric bsAb construct. Plasmids with high (N208D, E357Q, and S364K) and low (Q295E, L368D, K370S, N384D, Q418E, and N421D) pI mutations were obtained from Dr. Pamela Bjorkman (California Institute of Technology). Genes encoding the C1596 V_H_ and C_H_1 domains were subcloned into the high pI Fc plasmid, while the C952 variable light (V_L_) chain and constant lambda light chain genes were subcloned into the low pI Fc plasmid. The corresponding C952 V_H_ and C_H_1 domain genes were subcloned into an AbVec2.1 Fab expression plasmid; the C1596 V_L_ chain and constant kappa light chain genes were subcloned into an AbVec2.1 light chain expression plasmid. C1596-V_H_-high-pI-Fc, C1596 LC, C952-V_L_-low-pI-Fc, and C952 Fab plasmids were co-transfected in Expi293F cells using the ExpiFectamine transfection kit (ThermoFisher Scientific, A14525) in a 1:1.5:1:1.5 ratio; ExpiFectamine transfection enhancer was added 20 h post-transfection. CoV2-biRN1 was purified from cell supernatant via affinity chromatography using a HiTrap MabSelect SuRe (Cytiva) column. The eluted bsAb was concentrated using Amicon™ spin concentrators and then desalted on a HiPrep™ 26/10 Desalting column (Cytiva) against IEX Buffer A (50 mM Tris pH 8.5). Following desalting, the protein was concentrated and loaded onto a HiTrap Q HP column (Cytiva) and eluted with a 0-20% gradient of IEX Buffer B (50 mM Tris pH 8.5, 1M NaCl) over 30 column volumes (CV), followed by a 20-100% gradient of IEX Buffer B over 10 CVs. The desired protein product was identified via gel electrophoresis and further purified via size exclusion chromatography using either a HiLoad 16/600 Superdex 200 pg (Cytiva) or Superdex 200 Increase 10/300 GL (Cytiva) column against 1x TBS-Az. Pooled fractions corresponding to CoV2-biRN1 were stored in 1x TBS-Az at 4°C.

CoV2-biRN2 and CoV2-biRN3 were designed and cloned utilizing the dual variable domain immunoglobulin (DVD-Ig) bispecific format, as previously described (*85*). In brief, the VH and VL of either C952 or C1596 were covalently linked to the N-terminus of the IgG heavy chain and light chain, respectively, of the other antibody via a 24-mer (G_4_S)_4_G_4_ linker to create the outer variable domain of the second antibody. DVD-Ig constructs were co-transfected in Expi293F cells following a 1:1 ratio, with expression and purification following similar workflows as described above for monoclonal IgG antibodies. CoV2-biRNs 2 and 3 were stored in 1x TBS-Az at 4°C.

CoV2-biRNs 4 through 9 were designed and cloned utilizing a tandem single-chain fragment variable (scFv) bispecific format. Each arm of the tandem scFv bsAb consisted of an outer domain scFv linked to an inner domain scFv via a 25-mer (Gly_4_Ser)_5_ linker. Tandem scFvs were fused to the human immunoglobulin Fc C_H_2-C_H_3 domains via a human IgG hinge domain. CoV2-biRNs 4, 5, 8, and 9 utilized the IgG1 hinge (EPKSCDKTHTCPPCP), whereas CoV2-biRNs 6 and 7 used the IgG3 hinge (ELKTPLGDTTHTCPRCP(EPKSCDTPPPCPRCP)_3_).

Constructs were cloned into an AbVec2.1 expression vector and transfected in Expi293F cells following the same expression and purification workflow described above for monoclonal IgG antibodies. CoV2biRNs 4 through 9 were stored in 1x TBS-Az at 4°C. See Data Table 1 and Supplemental Figure 7 for CoV2-biRN bsAb construct design and purification representations.

### Protein cloning, expression, and purification

All recombinant viral proteins were cloned into a mammalian expression vector with an N-terminal signal peptide and a C-terminal AviTag followed by a polyhistidine tag. Constructs were transfected in Expi293F cells and expressed for four days at 37°C. For biotinylated constructs, cells were co-transfected with a plasmid encoding for the *Escherichia coli* biotin ligase (BirA) obtained from the laboratory of Dr. Kai Zinn (California Institute of Technology). Recombinant proteins were isolated from cellular supernatants and purified via affinity chromatography using a HisTrap HP column (Cytiva) and size-exclusion chromatography on either HiLoad 16/600 Superdex 200 pg (Cytiva) or Superdex 200 Increase 10/300 GL (Cytiva) columns using an AKTA pure 25 M1 system (Cytiva). All proteins were stored in 1x TBS-Az at 4°C. Detailed sequence and design information for all recombinant viral protein constructs used in this study can be found in Data Table 2. pcDNA3-sACE2(WT)-8his encoding soluble human ACE2 (UniProt Q9BYF1, residues 1-732) was a gift from Erik Procko (Addgene plasmid #149268) and was expressed and purified as described above for the other recombinant proteins.

### Enzyme-linked immunosorbent assays

ELISAs to measure binding between monoclonal Abs and viral antigens were performed by immobilizing viral antigens directly on high-binding 96-well assay plates (Corning, 9018) at a concentration of 2 µg/ml in 1x TBS-Az for 20 h at 4°C. After overnight incubation, plates were washed with washing buffer (1x TBS + 0.05% Tween-20; TBS-T), followed by incubation for 1 h at RT in blocking buffer (TBS + 0.05% Tween-20 + 1% non-fat dry milk + 1% goat serum; TBS-TMS). After washing the plates with TBS-T washing buffer, serial dilutions of monoclonal antibodies were added to the plates at a top concentration of 100 µg/mL in TBS-TMS blocking buffer and incubated for 2 h at RT. Plates were washed in TBS-T and incubated with horseradish peroxidase (HRP)-conjugated goat anti-human Ig Fc antibody (Southern Biotech, 2047-050) at a dilution of 1:4000 in TBS-TMS for 30 min at RT. After washing the plates with TBS-T, 100 µl of 1-Step™ Ultra TMB-ELISA substrate (Thermo Scientific, 34029) was added to each well for approximately 2-3 min. Reactions were quenched with 100 µl per well of a 1 M HCl solution, and absorbance at 450 nm was measured with a Tecan Infinite^®^ M Plex microplate reader using iControl 2.0 software. Data obtained were plotted with GraphPad Prism 10.

### Biolayer interferometry

BLI measurements were performed on an Octet Red96 system (FortéBio) at 20°C in Octet Buffer (TBS + 0.1% Bovine Serum Albumin + 0.02% Tween-20). For assays used to calculate binding kinetic data, IgGs or Spike trimers were directly immobilized onto ProA or streptavidin (SA) biosensors (Sartorius) and dipped into a concentration series of monomeric analytes (top concentrations ranging from 1 µM to 10 µM) for 120s and allowed to dissociate in Octet buffer for 300-600 s. For assays using ProA or HIS1K biosensors (Sartorius), regeneration of biosensors was achieved using 10 mM Glycine, pH 1.5, and neutralized with Octet Buffer. Kinetic analyses were used after the subtraction of reference curves to derive on/off rates (*k*_a_/*k*_d_) and binding constants (*K*_D_s) using a 1:1 binding model in the ForteBio Data Analysis 9.0 software and GraphPad Prism 10. Reported affinities represent the average of two independent experiments.

BLI experiments that were not used to derive binding affinities or kinetic constants were done using a single high concentration to qualitatively distinguish binding versus no binding.

### SARS-CoV-2 pseudotyped reporter viruses

The plasmids and protocol for the generation of pseudotyped virus were as previously described, with minor modifications (*77*). Plasmids used included pHAGE-CMV-Luc2-IRES-ZsGreen-W, HDM-Hgpm2, HDM-tat1b, pRC-CMV-Rev1b, and HDM-SARS-CoV-2-Spike-del21. A panel of plasmids encoding for spike proteins from the Omicron VOCs was based on SARS-CoV-2 (Genbank MN985325.1) with a D614G mutation and a 21-residue truncation at the C-terminus. Sequence details for Omicron VOC Spike plasmids can be found in Data Table 2. Briefly, HEK293T cells were transfected with plasmids using FuGENE^®^ HD (Promega). At 72 h post-transfection, supernatants containing generated pseudovirions were harvested via centrifugation at 300 x g, filtered through a 25 mm syringe filter with 0.45 µm Supor membrane (Pall, 4614), and stored at -80°C. Generated pseudotyped virus stocks were titrated on HeLa-hACE2-TMPRSS2 cells to determine infectivity.

### SARS-CoV-2 pseudotyped virus neutralization assay

*In vitro* pseudovirus neutralization assays were performed as previously described, with minor modifications (*77*). Briefly, 8×10^3^ HeLa-hACE2-TMPRSS2 cells were plated in 100 µl of D10 media per well in a 96-well plate (Fisher Scientific, cat no. 12-565-383). Plated cells were incubated at 37°C and 5% CO_2_. Eighteen to twenty hours later, serial dilutions of antibody inhibitors were prepared at a top concentration of 100 µg/mL and incubated with a 1:1 volume of titrated SARS-CoV-2 pseudovirus at 37°C and 5% CO_2_ for 1 h. After the incubation, 50 µl of D10 media was removed from the plated HeLa-hACE2-TMPRSS2 cells and replaced with 100 µl of the antibody-pseudovirus mixture. Plated cells with inhibitor and pseudovirus were incubated at 37°C and 5% CO_2_. Forty-eight hours later, 100 µl was removed from the plated cells and a 1:1 volume of Britelite™ Plus to remaining media was added to the plated cells. Microplates were shaken for 30 seconds in an orbital manner and luminescence was analyzed with no attenuation and 1 s integration time using the Tecan Infinite^®^ M Plex microplate reader and iControl 2.0 software. Data were plotted with GraphPad Prism 10. Reported IC_50_ values represent the average of three independent experiments.

### Mouse experiments

All experiments in this study involving animals were conducted in strict compliance with the United States Animal Welfare Act and following the National Institutes of Health Guide for the Care and Use of Laboratory Animals. The research protocols were approved by the Institutional Animal Care and Use Committee at The Rockefeller University (21042-H). K18-hACE2 transgenic mice [B6.Cg-Tg(K18-ACE2)2Prlmn/J, 034860] (*74, 75*) were obtained from the Jackson Laboratory and housed in a standard BSL1 facility at the Rockefeller Comparative Biosciences Center with ad libitum access to food and water under a 12 h-12 h light-dark cycle. K18-hACE2 mice (n = 5 per group; 8 weeks old) were injected intraperitoneally with the indicated inhibitor at a dose of 5 mg kg^-1^ either twenty-four hours before (prophylaxis arm) or 4 hours after (treatment arm) study day 0. On day 0, all mice were inoculated with 2 x 10^4^ PFU of SARS-CoV-2 XBB.1.5 via intranasal administration. At 3 days after infection, the left lung was homogenized in Trizol LS reagent (Ambion, Life Technologies), and total RNA was extracted by phase separation using chloroform (Sigma-Aldrich). RNA in the aqueous phase was precipitated using isopropanol (Sigma-Aldrich). Pelleted RNA was then washed with 75% ethanol and resuspended in nuclease-free water. Lung viral load was quantified by qRT-PCR using PowerSYBR Green RNA-to-CT, 1-Step Kit (ThermoFisher Scientific) and StepOne Plus Real-Time PCR system (Applied Biosystems). The primers (2019-nCoV_N1-F: 5′-GACCCCAAAATCAGCGAAAT-3′ and 2019-nCoV_ N1-R: 5′-TCTGGTTACTGCCAGTTGAATCTG-3′) targeted RNA sequences that encode the nucleocapsid protein of SARS-CoV-2 XBB.1.5. A standard, 2019-nCoV_N_Positive Control 10006625 was obtained from IDT.

### Pseudovirus-based full spike deep mutational scanning

The XBB.1.5 full spike deep mutation scanning libraries were developed as previously described (*88*). To map escape mutations for the C952-C1596 IgG cocktail and CoV2-biRN5 bsAb, ∼1 million infectious units of library virus were mixed with the antibody cocktail or bsAb at three increasing concentrations; for the antibody cocktail these were equivalent to IC_99_, 3xIC_99_, and 9xIC_99_ and for the bsAb these were equivalent to IC_99_, 4xIC_99_, and 16xIC_99_ concentrations as determined by pseudovirus neutralization assays performed on HEK-293T-medium-ACE2 cells (*93*). The library virus was incubated with the antibodies for 45 min at 37°C before infecting HEK-293T-medium-ACE2 cells. Fifteen hours after infection, the viral genomes were collected from the cells, followed by barcode sequencing, as previously described(*87*). Two biological replicates (using independent deep mutational scanning libraries) were used to map escape for each antibody. A VSV-G pseudotyped virus was used as a non-neutralized standard to determine escape, as previously described (*87*). Analysis of the experimental data was performed employing a biophysical model described previously (*94*) and implemented in a polyclonal package v6.9 (https://jbloomlab.github.io/polyclonal/). The complete analysis pipeline for the C952-C1596 antibody cocktail and the CoV2-biRN5 bsAb and the underlying data can be accessed at https://github.com/dms-vep/SARS-CoV-2_XBB.1.5_spike_DMS_Barnes_mAbs. Interactive plots showing log2 IC_90_ fold change in neutralization for the cocktail and the bsAb can be found at https://dms-vep.org/SARS-CoV-2_XBB.1.5_spike_DMS_Barnes_mAbs/htmls/cocktail-C952-C1596_mut_icXX.html and https://dms-vep.org/SARS-CoV-2_XBB.1.5_spike_DMS_Barnes_mAbs/htmls/CoV-biRN-10_mut_icXX.html, respectively.

### Cryo-EM sample preparation

Quantifoil Cu 1.2/1.3 300 mesh grids were glow-discharged for 60 s at 10 mA (PELCO easiGlow) before sample application. For Spike-Fab complexes, 3 mg/mL purified Spike 6P was incubated with antibody Fab fragments at a 1:1.1 spike protomer to Fab molar ratio for 30 min in 1x TBS. Immediately prior to depositing 3 µL of the complex onto grids, fluorinated octyl-maltoside (FOM, Anatrace) was added to the sample to a final concentration of 0.02% w/v. Grids were blotted for 3 s at room temperature, 100% humidity, blot force 0, plunge-frozen in ethane, and stored in liquid nitrogen until data collection.

### Cryo-EM data collection and processing

Single-particle cryo-EM datasets were collected on a Titan Krios transmission electron microscope (Thermo Fisher) equipped with a K3 direct electron detector (Gatan), operating at 300 kV for all Spike-Fab complexes. Movies were acquired in an automated fashion using SerialEM v3.8 and a 3 x 3 multishot data acquisition pattern, with a total dose of 60 e^−^/Å^2^ accumulated across 40 frames with a calibrated pixel size of 0.8521 Å (C1533-S 6P and C1596-S 6P) or 0.8677 Å (C952-S 6P) and a defocus range of -1 to -2.5 µm.

For all datasets, motion correction, CTF estimation, reference-free particle-picking and extraction were carried out using cryoSPARC v4.1 (*95*). Micrographs displaying ice contamination, CTF estimated resolutions worse than 5 Å and ice thickness greater than a score of 1.25 were discarded. A subset of 4x-downsampled reference-free particles were used to generate ab intio reconstructions in cryoSPARC, followed by heterogenous refinement of the entire dataset to generate an initial reference model. Particles corresponding to Spike-Fab complexes were iteratively refined using 2D and 3D classification in cryoSPARC and/or Relion (*96*), followed by re-extraction without binning and non-uniform refinement in cryoSPARC. For local refinement of the C1533 Fab V_H_-V_L_ – NTD interface, 3D classification in Relion (k=4) yielded 559,601 particles, which were refined with a mask and C1 symmetry in cryoSPARC. Resolution of all the maps were estimated according to the 0.143 Fourier shell correlation (FSC) cut-off criteria in cryoSPARC. Further details of data collection and processing workflows for all datasets can be found in Supplemental Table 3 and Supplemental Figures 1, 3, and 7.

### Cryo-EM structure modeling, refinement, and analyses

Global non-uniform maps (C1596-S 6P & C952-S 6P) or the locally refined map (C1533 V_H_V_L_-NTD) were used for model building, with both sharpened and unsharpened maps used for model refinement (unsharpened maps used for glycan refinement). Initial coordinates for each complex were obtained by docking individual chains from homology models (Supplemental Table 3) into cryo-EM maps using UCSF ChimeraX (*97*). Sequences were manually updated in Coot (*98*) and refined using iterative rounds of refinement in Phenix (*99*) and manual building in Coot. Glycans were modeled at potential *N*-linked glycosylation sites using the carbohydrate module in Coot. Validation of refined model coordinates was performed in MolProbity (*100*).

UCSF ChimeraX v1.7 was used for structure visualization. Buried surface areas reported in Supplemental Table 2 and epitope assignment were calculated using PDBePISA (*101*) and a 1.4 Å probe. Potential hydrogen bond (less than 4 Å and with a A-D-H angle greater than 90°) and van der Waals interaction assignments were performed using ChimeraX.

### Quantification and Statistical Analysis

The p-values in Figure 6 were determined through one-way ANOVA, correcting for multiple comparisons with Dunnett’s test with comparison to the untreated control. The levels of significance are indicated as follows: **p<0.01; *p<0.05. The EC_50_, IC_50_, and IC_90_ values were calculated by fitting the data to dose-response curves using GraphPad Prism 10. The EC_50_ values were determined using a [Agonist] vs response – Variable slope (four parameters) model. The IC_50_ values were determined using a [Inhibitor] vs response – Variable slope (four parameters) model. The IC_90_ values were determined using a [Agonist] vs response – Find ECanything model with F = 10. BLI data were processed using the ForteBio Data Analysis 9.0 software. Data points were subtracted from data from reference wells, aligned to the baseline step for the Y-axis and inter-step correction was applied, aligning to the dissociation phase. Finally, data were processed using Savitzky-Golay filtering. Processed data were plotted and processed with GraphPad Prism 10. K_D_ values were calculated by fitting data points to a nonlinear regression curve fit using an Association then Dissociation model, accounting for a 1:1 binding of ligand to analyte.

## Acknowledgments

We thank members from the laboratory of Dr. Pamela Bjorkman (California Institute of Technology) and Dr. Peter S. Kim (Stanford) for cell lines, SARS-CoV-2 spike expression constructs, and technical assistance with pseudovirus neutralization assays. Cryo-EM data for this work was collected at the Stanford-SLAC cryo-EM resource center with support from Dr. Elizabeth Montabana. Some of this work was performed at the Stanford-SLAC Cryo-EM Center (S2C2), which is supported by the National Institutes of Health Common Fund Transformative High-Resolution Cryo-Electron Microscopy program (U24 GM129541). The content is solely the responsibility of the authors and does not necessarily represent the official views of the National Institutes of Health. The authors would also like to thank S2C2 personnel, Dr. Chensong Zhang and Dr. Patrick Mitchell, for their invaluable support and assistance. J.D.B., M.C.N., and P.D.B. are Howard Hughes Medical Institute (HHMI) Investigators. This article is subject to HHMI’s Open Access to Publications Policy. HHMI lab heads have previously granted a non-exclusive CC BY 4.0 license to the public and a sublicensable license to HHMI in their research articles. Pursuant to those licenses, the author-accepted manuscript of this article can be made freely available under a CC BY 4.0 license immediately upon publication. Finally, we thank all members of the Barnes lab for their careful review and feedback on the manuscript.

## Funding

This study was supported in part by funds from the Chan Zuckerberg Biohub (C.O.B), and Howard Hughes Medical Institute Emerging Pathogens Initiative (C.O.B). Additionally, C.O.B. is supported by the Howard Hughes Medical Institute Hanna Gray Fellowship, Rita Allen Foundation, Pew Biomedical Scholars Program, and is a Chan Zuckerberg Biohub investigator.

This work was supported in part by NIH/NIAID grant R01 AI141707 to J.D.B. A.A.R. is supported by the National Science Foundation Graduate Research Fellowship Program and the National Institutes of Health Training Grant (T32 AI00729037). V.A.B. is supported by the Boehringer Ingelheim Fonds PhD fellowship. Z.W. received support from the SNF Institute for Global Infectious Disease for the Advancement of Translational Research, partially funded by grant # UL1 TR001866 from the National Center for Advancing Translational Sciences (NCATS), under the National Institutes of Health (NIH) Clinical and Translational Science Award (CTSA) program. J.D.B, P.D.B, and M.C.N are Howard Hughes Medical Institute investigators.

## Author contributions

A.A.R. and C.O.B. conceived, designed, and analyzed the experiments. A.A.R. performed and analyzed all experiments with assistance from others: M.P., Z.W., M.E.A., Y.E.L., M.R.E., J.P., I.R., and T.C. supported cloning, expression, and purification of recombinant protein constructs; M.P helped perform binding experiments; M.P., Y.E.L., and J.P. helped perform pseudovirus neutralization assays; M.E.A., G.E.N., and C.O.B. helped perform and analyze cryo-EM structural data; B.D. and J.D.B. performed and analyzed pseudovirus-based deep mutational scanning assays; V.A.B., P.D.B., and M.C.N. performed and analyzed challenge experiments. The paper was written by A.A.R. and C.O.B. with assistance from all co-authors.

## Competing interests

The Rockefeller University has filed a provisional patent application in connection with monoclonal antibodies described in this work on which Z.W. and M.C.N. are inventors (US patent 17/575,246). J.D.B. is on the scientific advisory boards of Invivyd, Aerium Therapeutics, Apriori Bio, and the Vaccine Company. J.D.B. consults for GSK and Moderna. B.D. consults for Moderna. J.D.B. and B.D. are inventors on Fred Hutch licensed patents related to viral deep mutational scanning.

## Data and materials availability

The atomic models and cryo-EM maps generated for the C1533 V_H_V_L_-NTD (local refinement), C1596-S 6P, and C952-S 6P complexes have been deposited to the Protein Data Bank (PDB; http://www.rcsb.org/) and the Electron Microscopy Databank (EMDB; http://www.emdataresource.org/) under accession codes PDB: 9BJ2, 9BJ3, and 9BJ4 and EMD-44627, -44628, and -44629, respectively.

The code and pipeline used to analyze the deep mutational scanning data is available at https://github.com/dms-vep/SARS-CoV-2_XBB.1.5_spike_DMS_Barnes_mAbs. Interactive visualization of the escape results is available at https://dms-vep.org/SARS-CoV-2_XBB.1.5_spike_DMS_Barnes_mAbs/htmls/cocktail-C952-C1596_mut_icXX.html and https://dms-vep.org/SARS-CoV-2_XBB.1.5_spike_DMS_Barnes_mAbs/htmls/CoV-biRN-10_mut_icXX.html.

## Supplemental Materials

**Supplemental Figure 1.**
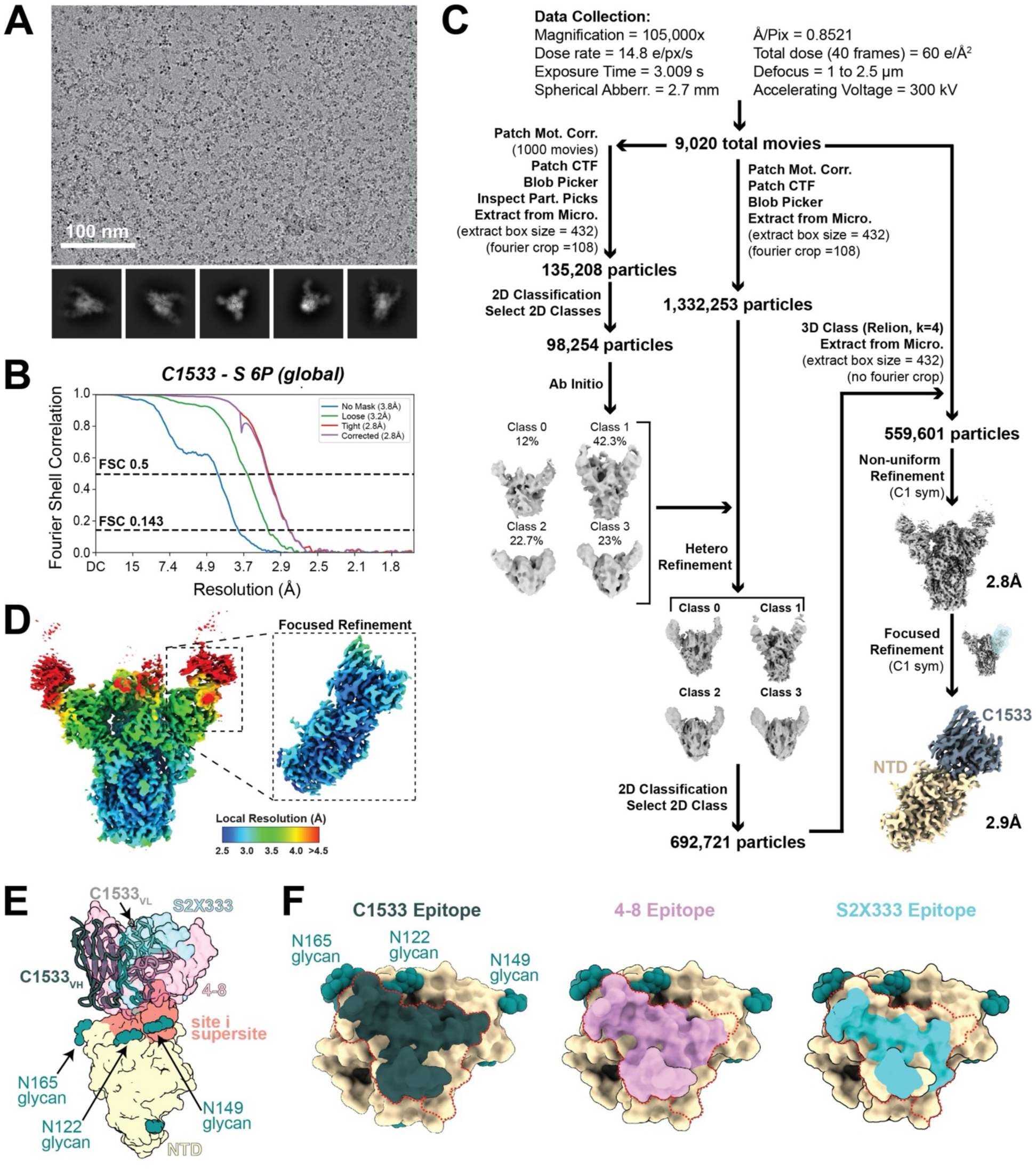
C1533 Fab in complex with SARS-CoV-2 S 6P. Related to Figure 1. **(A)** Representative micrograph and 2Dclass averages selected from the total dataset of C1533-S 6P. **(B)** Gold-standard FSC plots for the C1533-S 6P global refinement. **(C)** Data collection and processing workflow. Final focused-refined map for C1533 variable domains (slate gray) bound to the NTD subunit (wheat) to 2.9Å resolution is shown. **(D)** Local resolution estimations calculated in cryoSPARC for the C1533-S 6P global refinement and C1533_VhVl_-NTD focused refinement (inset). **(E)** Superimposition of 4-8 (PDB: 8DLR, pink), S2X333 (PDB: 7LXW, cyan), and C1533 V_H_ (dark slate _gr_ay) and V_L_ (light gray) domains onto the SARS-CoV-2 Gamma P.1 NTD (PDB: 8DLR, wheat) after alignment on NTD residues 14-19, 67-69, 77-85, 100-122, 126-163, and 236-251 for a composite figure. **(F)** Surface rendering of SARS-CoV-2 Gamma P.1 NTD (PDB: 8DLR, wheat) with epitope footprints for C1533, 4-8, and S2X333 depicted, relative to the antigenic supersite (outlined in red).

**Supplemental Figure 2.**
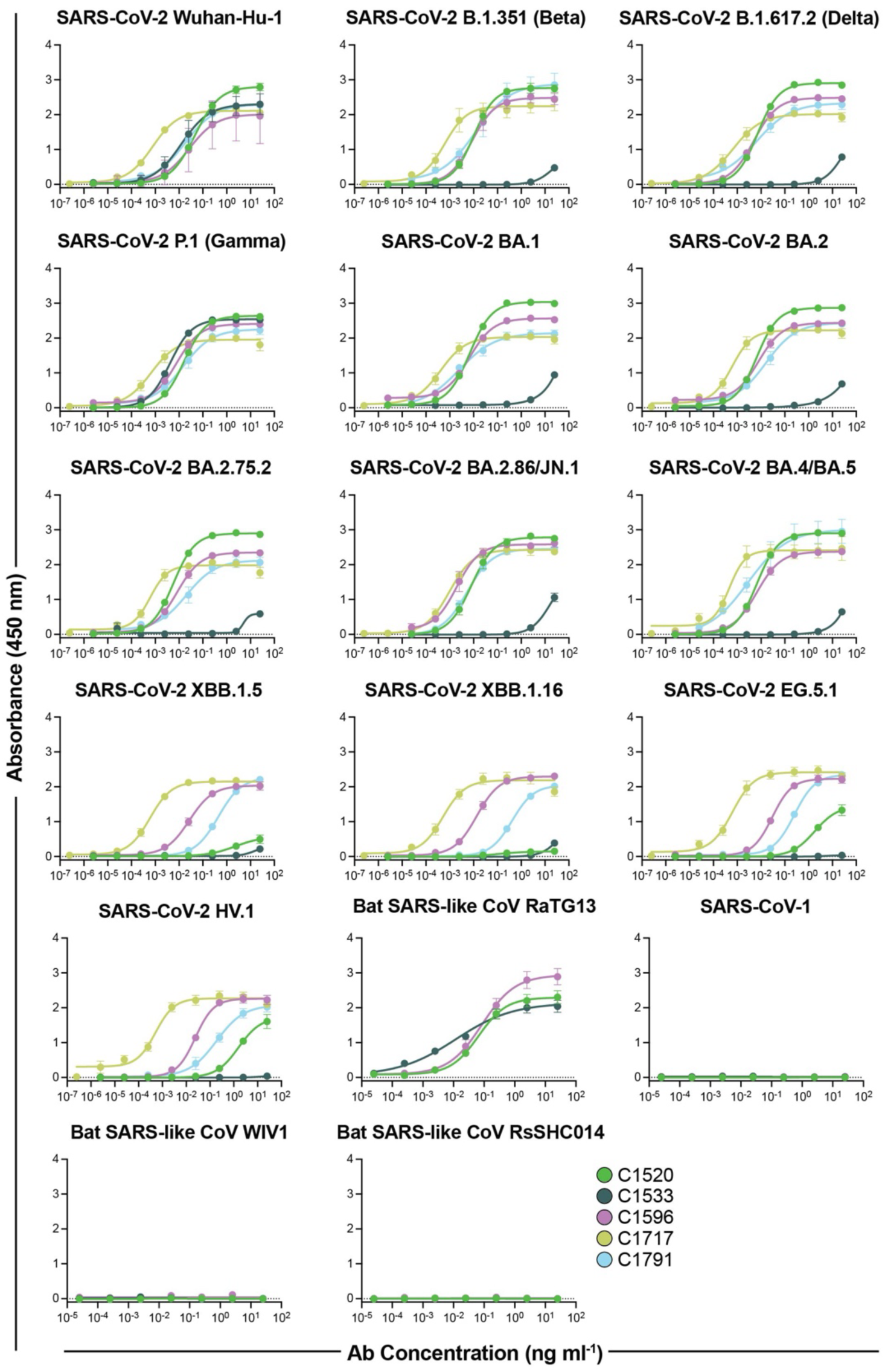
ELISA curves for NTD-specific IgG antibodies binding to SARS-CoV-2 VOC and SARS-like NTD proteins. Related to Figure 1. ELISAs to compare C1520 (green), C1533 (dark slate gray), C1596 (magenta), C1717 (lime), and C1791 (cyan) binding to directly-coated monomeric NTD protein constructs. Values represent the mean and standard error of the mean of three biological replicates (n=3).

**Supplemental Figure 3.**
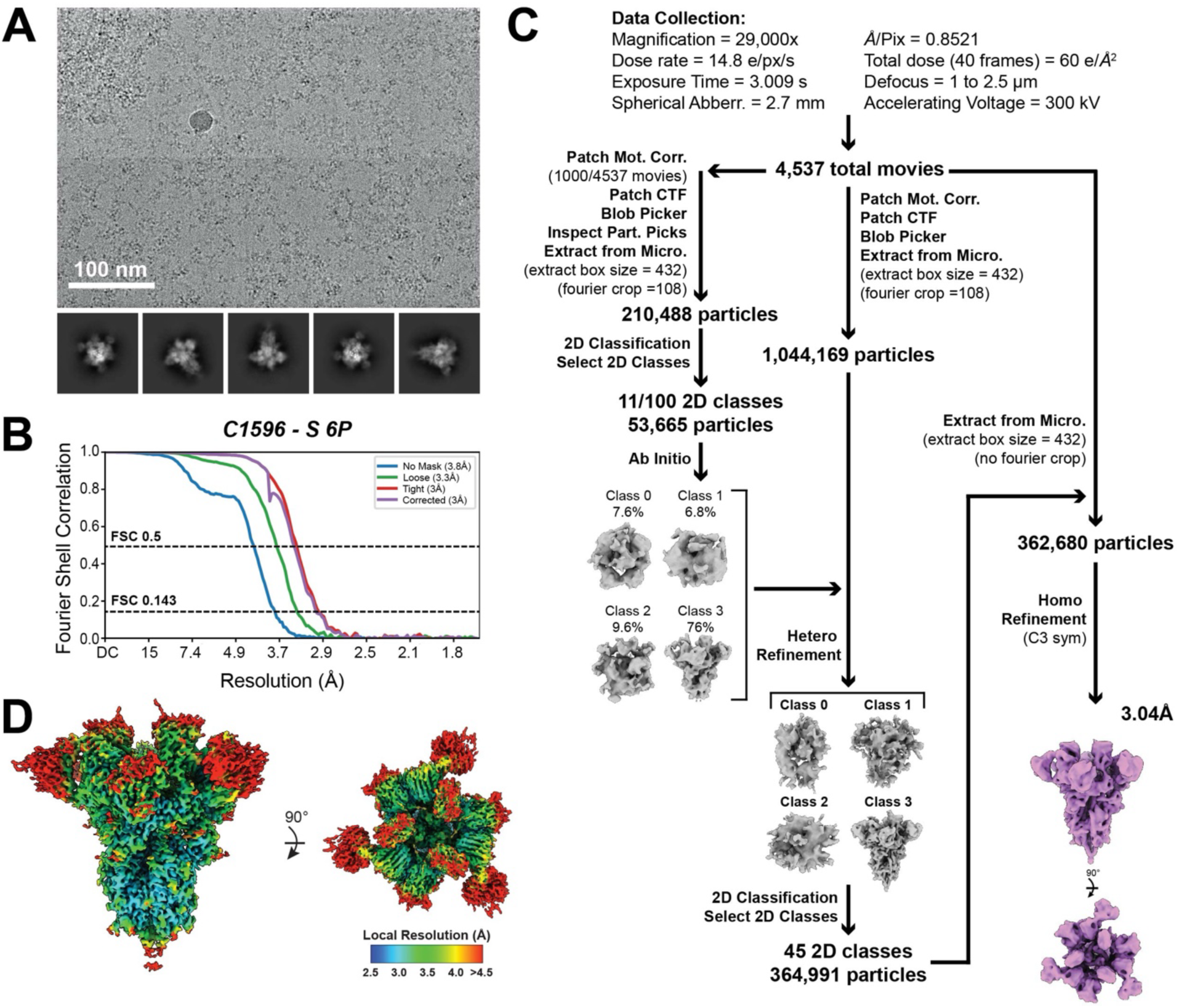
Cryo-EM data collection and processing workflow for C1596-SARS-CoV-2 S 6P complex. Related to Figure 2. **(A)** Representative micrograph and 2D class averages selected from the total dataset for C1596-S 6P. **(B)** Gold-standard FSC plots for C1596-S 6P. **(C)** Processing workflow for C1596-S 6P dataset in CryoSPARC. **(D)** Local resolution estimations for C1596-S 6P.

**Supplemental Figure 4.**
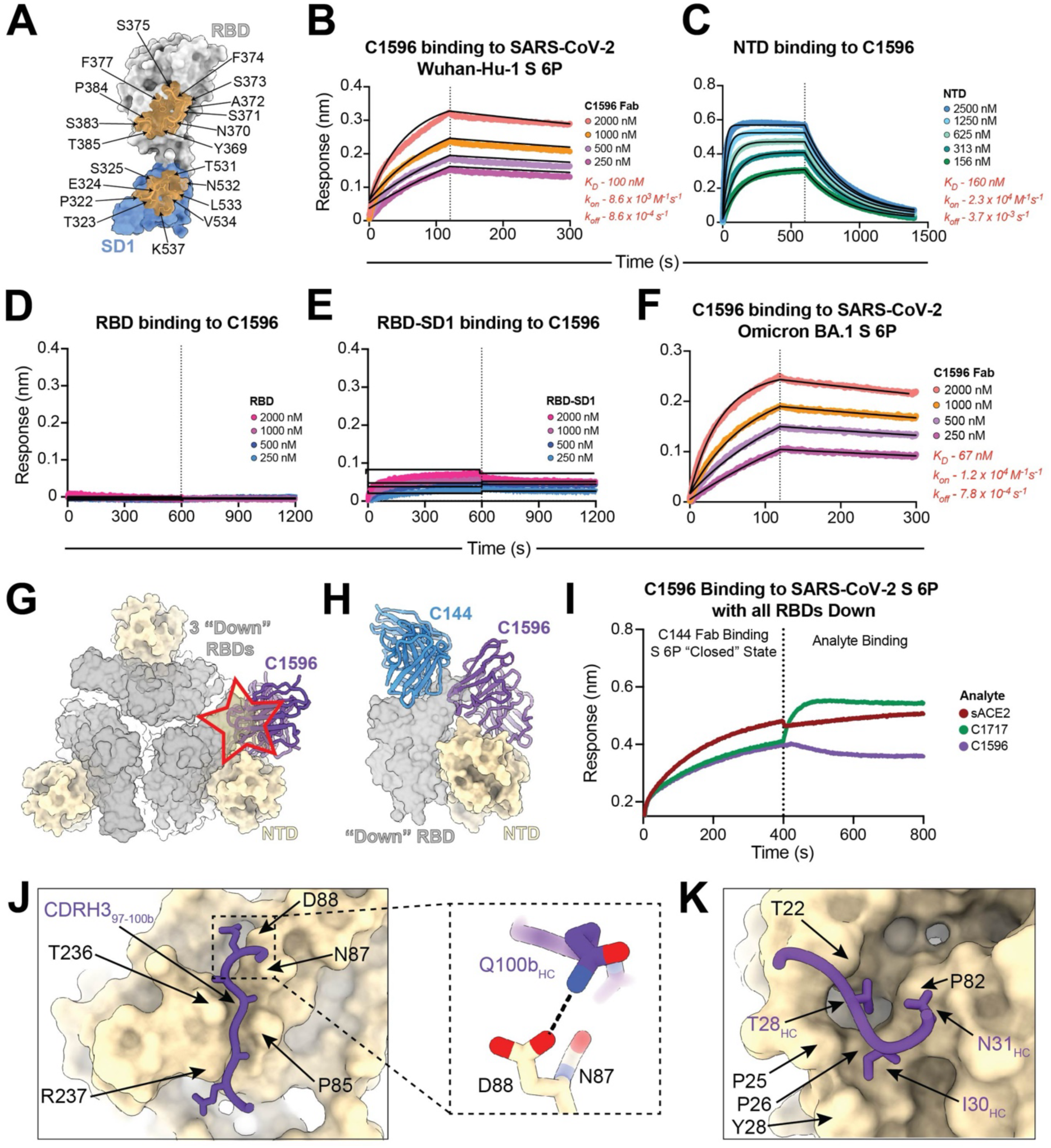
Characterization of C1596 molecular interactions with spike S1 domains. Related to Figure 2. **(A)** C1596 epitope footprint (brown) outlined on the RBD (light gray) and SD1 (cornflower blue) domains, with interfacing amino acids denoted. **(B-F)** BLI binding kinetics data of C1596 Fab binding to immobilized SARS-CoV-2 Wuhan-Hu-1 S 6P trimer **(B)** or SARS-CoV-2 BA.1 S 6P trimer **(F)**, and monomeric NTD **(C)**, monomeric RBD **(D)**, or monomeric RBD-SD1 **(E)** binding to immobilized C1596 IgG. The concentrations of analyte are indicated on each panel. The vertical dotted lines indicate the transition between association and dissociation phases. Fit curves are depicted as black solid lines. A K_D_ could not be determined in **(D)** or **(E)** due to the weak responses observed. **(G)** Modeling of C1596 V_H_-V_L_ (purple) on the SARS-CoV-2 S 6P trimer (PDB: 7K90) in the “closed” state, with all three RBDs (light gray) in the down position. Steric clash of C1596 with an adjacent down RBD is depicted as a red and yellow star. **(H)** Modeling of C1596 V_H_-V_L_ (purple) binding to the NTD (wheat) relative to C144 (blue) bound to an adjacent down RBD (PDB: 7K90), illustrating the lack of a clash between the two antibodies. **(I)** BLI experiment evaluating the ability of C1596 to bind S 6P when locked in the “closed” state. C144 Fab was bound to S 6P to conformationally lock the trimer in the “closed state”, followed by a second association with C1596 Fab. Inclusion of ACE2 is depicted to confirm the “closed” state. Association with C1717 is depicted to illustrate the relative shift of an NTD-binding antibody that is not dependent on the RBD conformation. The vertical dotted line represents the transition between the two association steps. **(J)** Representation of C1596 CDRH3 residues contacting a groove in the NTD. Inset: Stabilizing contact with residues Q100b of the C1596 CDRH3 mediated by a potential hydrogen bond with the NTD. **(K)** Representation of C1596 heavy chain engaging a hydrophobic pocket on the NTD utilizing framework region 1 and CDRH1 residues.

**Supplemental Figure 5.**
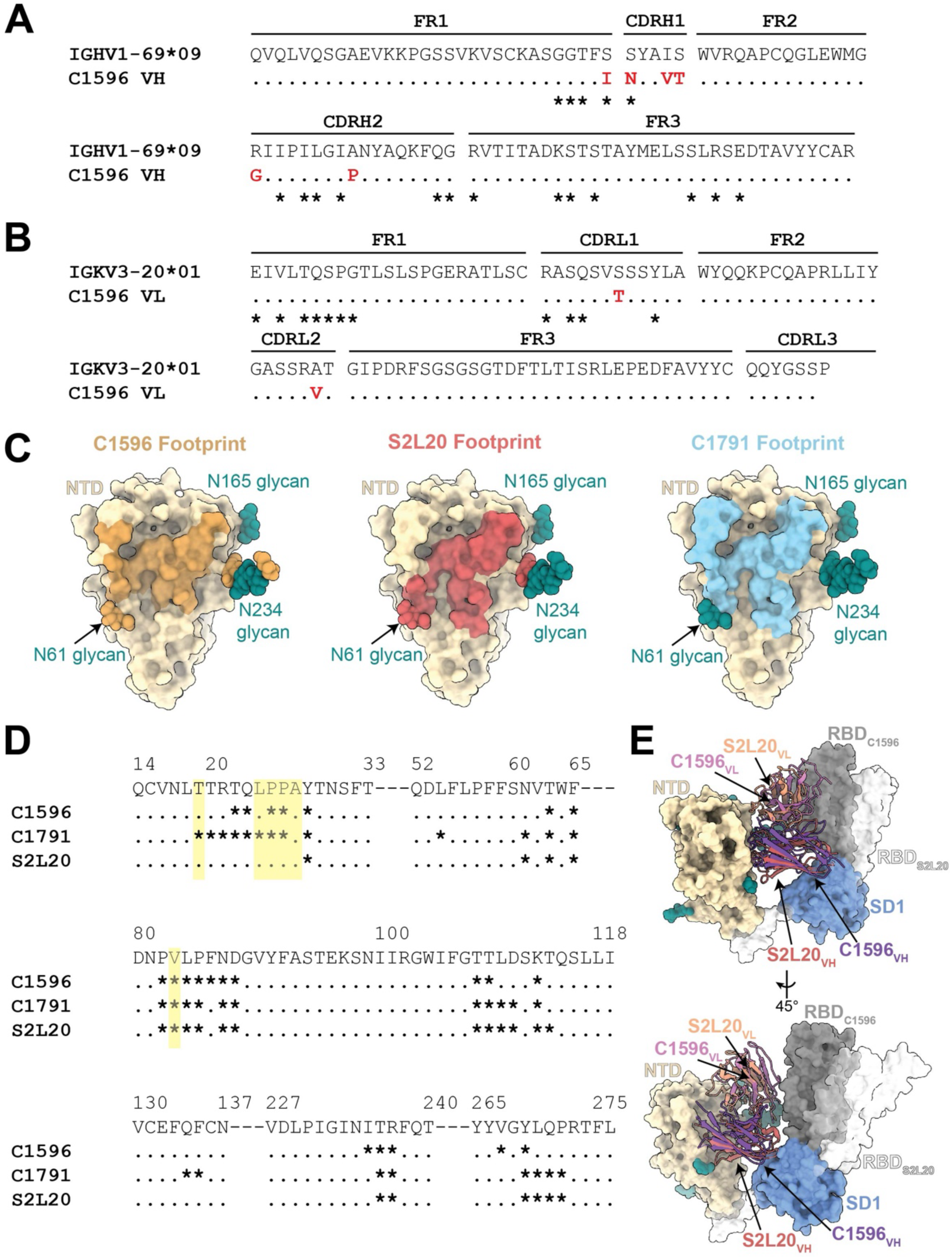
C1596 binds the glycopeptidic NTD epitope shared by C1791 and S2L20. Related to Figure 2. **(A-B)** Amino acid sequence alignment of C1596 **(A)** V_H_ and **(B)** V_L_ to germline reference sequences determined by V-Quest/IMGT. Mutations from the germline sequence are shown in red. Paratope residues of C1596 at the spike trimer interface are denoted with an asterisk. Framework regions, FR; and complementarity determining regions, CDRs, annotated using Kabat sequence numbering. **(C)** Surface rendering of SARS-CoV-2 Wuhan-Hu-1 NTD (wheat) with epitope footprints for C1596 (left, brown), S2L20 (middle, red), and C1791 (right, light blue) depicted. **(D)** Sequence representation of contacted NTD residues for C1596, C1791, and S2L20, depicted by asterisks. Sites where Omicron XBB.1.5 mutations overlap with NTD contacts are highlighted in ye_ll_ow. (_E)_ Overlay of V_H_ and V_L_ domains of C1596 (shades of purple) and S2L20 (PDB: 8GTQ; shades of red) after alignment on NTD residues 14-307, illustrating similar binding poses, but contrasting engagement with the RBD of the same protomer.

**Supplemental Figure 6.**
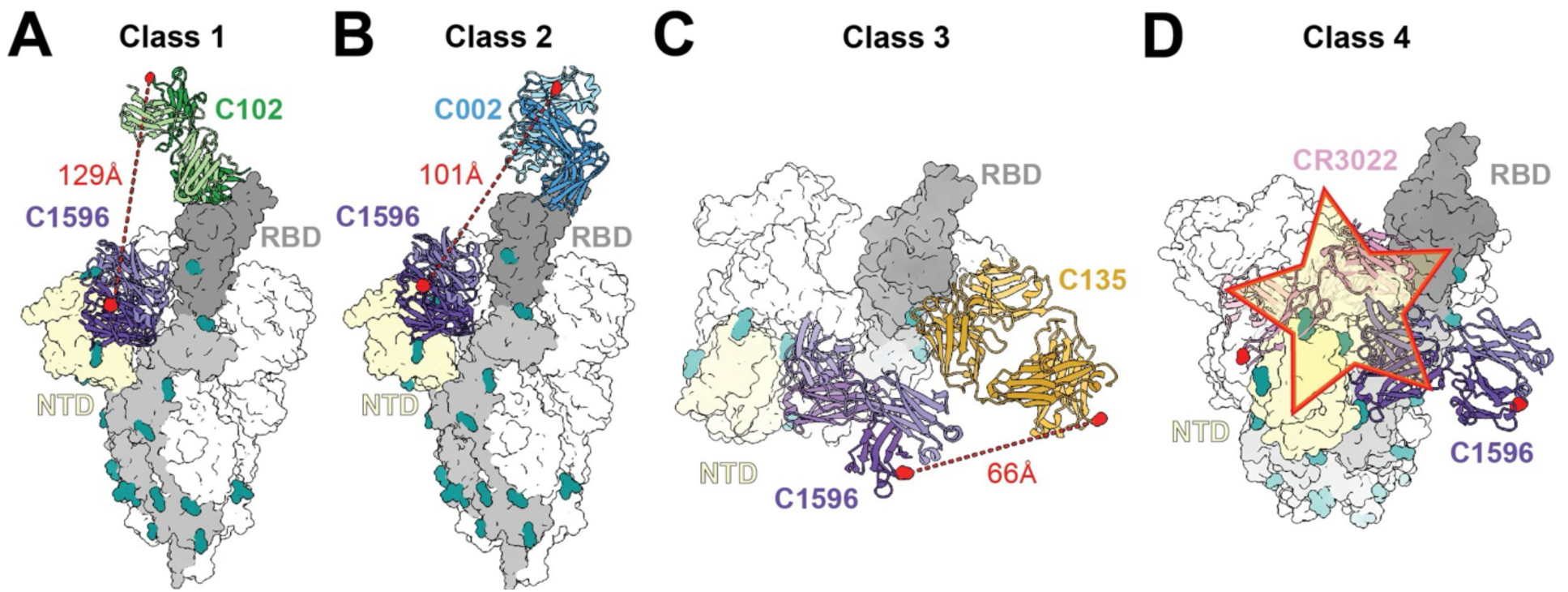
Structural modeling of distances between C1596 and RBD antibodies for bispecific design. Related to Figures 2 and 3. **(A-D)** Measurement of C⍺ distance (red dotted line) between the C termini of C1596 CH_1_ (purple) and the CH_1_ domains of **(A)** C102 (class 1 RBD epitope), **(B)** C002 (class 2 RBD epitope), **(C)** C135 (class 3 RBD epitope), and **(D)** CR3022 (class 4 RBD epitope). (A-D) A representative full-length Fab domain model was created for C1596 by aligning the variable heavy chain of C105 (PDB: 6XCA) to the variable heavy chain of C1596, aligning on residues 1-120 at the C⍺. NTD-C1596Fab models were then aligned to the NTD of the SARS-CoV-2 S glycoprotein (PDB: 7T67), aligning on NTD residues 14-307 at the C⍺. **(A)** The RBD-C102Fab model (PDB: 7K8M) was aligned to the RBD of the SARS-CoV-2 S, aligning on RBD residues 331-529 at the C⍺. **(B)** A representative full-length Fab domain model was created for C002by aligning the variable heavy chain of the C002 Fab (PDB: 7K8O) to the variable heavy chain of the C002 VH-VL model (PDB: 7K8T), as described for the C1596 Fab modeling. The RBD-C002Fab model was then aligned to the RBD of the SARS-CoV-2 S, as described in panel A. **(C)** A representative full-length Fab domain model was created for C135 by aligning the variable heavy chain of the C135 Fab (PDB: 7K8R) to the variable heavy chain of the C135 VH-VL model (PDB: 7K8Z), as described for C1596. The RBD-C135Fab model was then aligned to the RBD of the SARS-CoV-2 S, as described in panel A. **(D)** The RBD-CR3022 Fab model (PDB: 6YLA) was aligned to the RBD of the SARS-CoV-2 S model, as described in panel A.

**Supplemental Figure 7.**
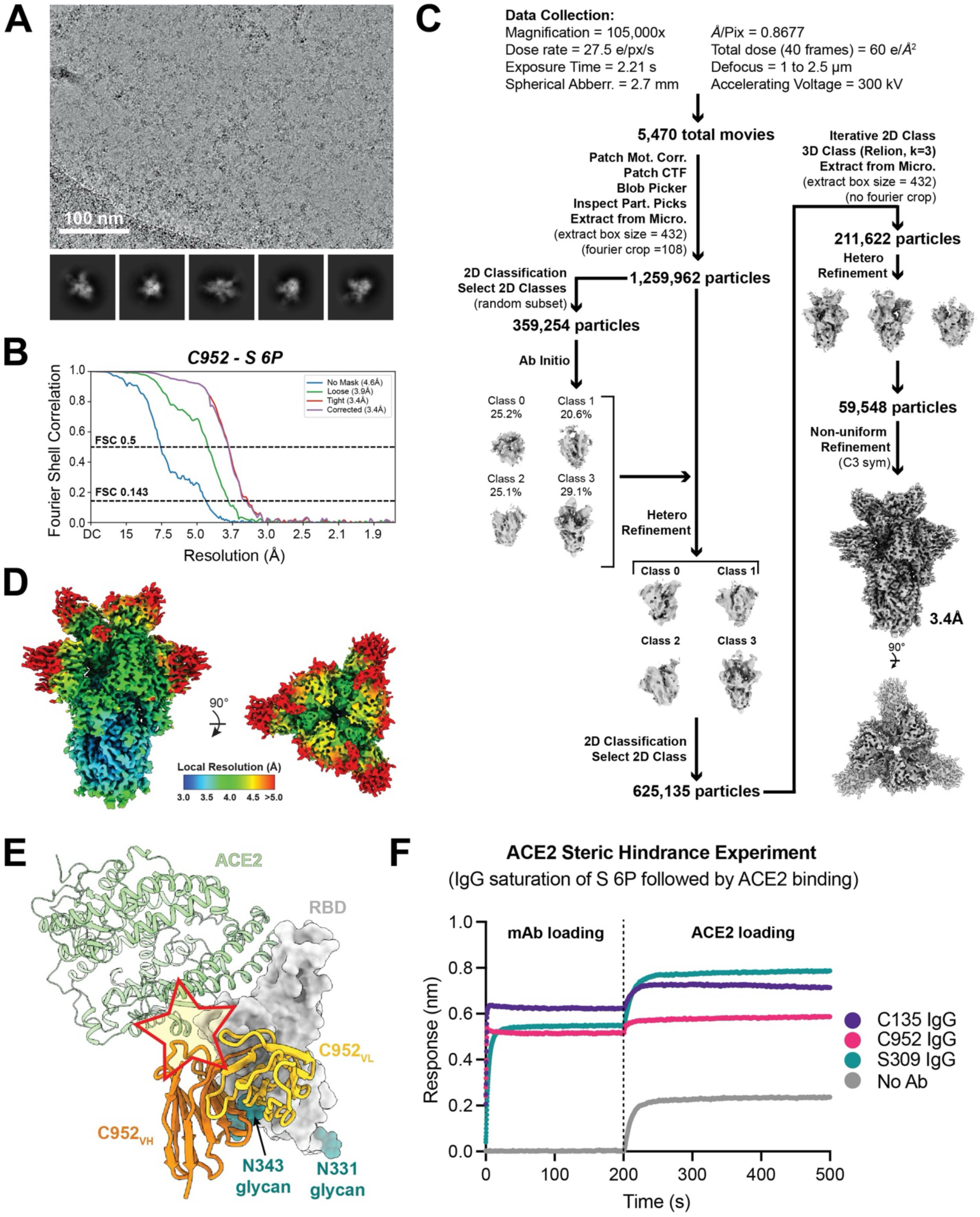
Cryo-EM data collection and data processing workflow for C952 Fab in complex with SARS-CoV-2 S 6P. Related to Figure 3. **(A)** Representative micrograph and 2D class averages selected from the total dataset of C952-S 6P. **(B)** Gold-standard FSC plots for the C952-S 6P global refinement. **(C)** Data collection and processing workflow. **(D)** Local resolution estimations calculated in cryoSPARC for the C952-S 6P global refinement. **(E)** Composite model of C952–RBD (shades of orange and gray, respectively) overlaid with soluble ACE2 (green; PDB 6M0J). Model was generated by aligning structures on 188 RBD Cα atoms. Potential clashes between ACE2 and C952 are highlighted by a yellow star. **(F)** mAb and ACE2 competition experiment by BLI. SARS-CoV-2 S 6P was immobilized on a streptavidin biosensor and saturated with either C135 (purple), C952 (pink), or S309 (green) mAb before dipping into a 1 µM solution of soluble ACE2 (as indicated by vertical dashed line). An ACE2 binding event (i.e., increase in y-axis response) indicates no competition for RBD binding between ACE2 and the corresponding mAb.

**Supplemental Figure 8.**
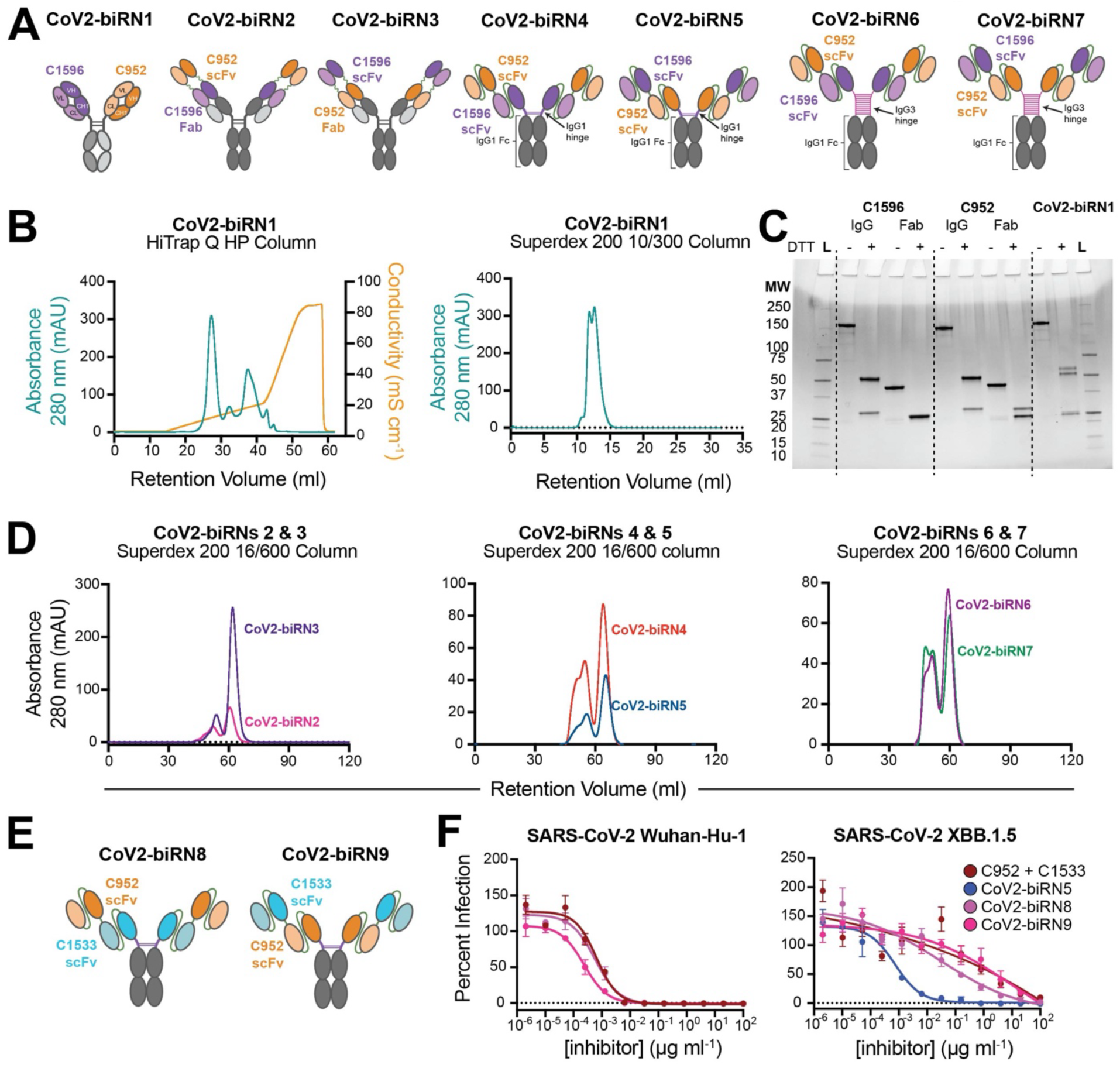
CoV2-biRN construct design and protein purification workflow. Related to Figure 4. **(A)** Schematic representation of CoV2-biRNs 1-7. **(B)** Representative chromatograph of (left) anion exchange followed by (right) size exclusion chromatography of CoV2-biRN1. **(C)** Protein gel of CoV2-biRN1 after anion exchange and size exclusion with parental antibodies ran as a reference under non-reducing or reducing conditions (L; ladder). **(D)** Representative chromatographs of the size exclusion purification of CoV2-biRNs 2 to 7. **(E)** Schematic of C952-C1533 tandem scFv bsAb constructs (CoV2-biRN8 and CoV2-biRN9). **(F)** Representative neutralization curves for CoV2-biRN8 and CoV2-biRN9 against (left) SARS-CoV-2 Wuhan-Hu-1 and (right) SARS-CoV-2 XBB.1.5 pseudoviruses. Data points represent the mean with standard error of the mean.

**Supplemental Table 1.** Binding kinetic data for monoclonal antibodies against SARS-CoV-2 VOC recombinant proteins. Related to Figures 1 and 3. N.B., no binding; N.T., not tested.

|  | C1520 |  |  | C1533 |  |  | C1596 |  |  |
| --- | --- | --- | --- | --- | --- | --- | --- | --- | --- |
| | $k_{on} (M^{-1}s^{-1})$ | $k_{off} (s^{-1})$ | $K_D (M)$ | $k_{on} (M^{-1}s^{-1})$ | $k_{off} (s^{-1})$ | $K_D (M)$ | $k_{on} (M^{-1}s^{-1})$ | $k_{off} (s^{-1})$ | $K_D (M)$ |
| SARS-CoV-2 D614G S 6P | *N.T.* | *N.T.* | *N.T.* | *N.T.* | *N.T.* | *N.T.* | 8.62E+03 | 8.58E-04 | 9.954E-08 |
| SARS-CoV-2 BA.1 S 6P | *N.T.* | *N.T.* | *N.T.* | *N.T.* | *N.T.* | *N.T.* | 1.17E+04 | 7.77E-04 | 6.66E-08 |
| Wuhan-Hu-1 NTD | 1.59E+04 | 3.93E-04 | 2.48E-08 | 3.20E+04 | 5.47E-04 | 1.71E-08 | 3.02E+04 | 3.79E-03 | 1.25E-07 |
| B.1.351 NTD | 2.05E+04 | 5.55E-04 | 2.71E-08 | *N.B.* | *N.B.* | *N.B.* | 2.45E+04 | 3.46E-03 | 1.41E-07 |
| B.1.617.2 NTD | 3.00E+04 | 1.41E-03 | 4.70E-08 | *N.B.* | *N.B.* | *N.B.* | 2.43E+04 | 3.00E-03 | 1.24E-07 |
| P.1 NTD | 1.31E+04 | 3.01E-03 | 2.30E-07 | 4.31E+04 | 1.07E-03 | 2.49E-08 | 2.35E+04 | 6.63E-03 | 2.82E-07 |
| BA.1 NTD | 3.55E+04 | 4.18E-03 | 1.18E-07 | *N.B.* | *N.B.* | *N.B.* | 4.47E+04 | 3.94E-03 | 8.82E-08 |
| BA.2 NTD | 3.60E+04 | 5.14E-04 | 1.43E-08 | *N.B.* | *N.B.* | *N.B.* | 4.03E+04 | 2.76E-02 | 6.85E-07 |
| BA.2.75.2 NTD | 3.07E+04 | 5.69E-03 | 1.85E-07 | *N.B.* | *N.B.* | *N.B.* | 3.63E+04 | 2.20E-02 | 6.07E-07 |
| BA.2.86/JN.1 NTD | 1.96E+04 | 1.10E-03 | 5.63E-08 | *N.B.* | *N.B.* | *N.B.* | 7.61E+04 | 5.55E-04 | 7.29E-09 |
| BA.4/BA.5 NTD | 4.63E+04 | 5.28E-04 | 1.14E-08 | *N.B.* | *N.B.* | *N.B.* | 7.27E+04 | 4.90E-02 | 6.74E-07 |
| XBB.1.5 NTD | *N.B.* | *N.B.* | *N.B.* | *N.B.* | *N.B.* | *N.B.* | 9.90E+04 | 5.24E-02 | 5.29E-07 |
| XBB.1.16 NTD | *N.B.* | *N.B.* | *N.B.* | *N.B.* | *N.B.* | *N.B.* | 8.92E+04 | 5.27E-02 | 5.91E-07 |
| EG.5.1 NTD | *N.B.* | *N.B.* | *N.B.* | *N.B.* | *N.B.* | *N.B.* | 1.08E+05 | 4.43E-02 | 4.10E-07 |
| HV.1 NTD | *N.B.* | *N.B.* | *N.B.* | *N.B.* | *N.B.* | *N.B.* | 7.07E+04 | 3.91E-02 | 5.53E-07 |
|  | C952 |  |  |  |  |  |  |  |  |
| | $k_{on} (M^{-1}s^{-1})$ | $k_{off} (s^{-1})$ | $K_D (M)$ | | | | | | |
| Wuhan-Hu-1 RBD | 2.78E+05 | 1.76E-03 | 6.35E-09 |  |  |  |  |  |  |
| BA.1 RBD | 3.01E+05 | 1.70E-04 | 6.27E-09 |  |  |  |  |  |  |
| BA.2.75.2 RBD | 2.93E+05 | 3.43E-04 | 1.73E-08 |  |  |  |  |  |  |
| BA.2.86 RBD | 1.41E+02 | 1.13E-01 | 8.00E-04 |  |  |  |  |  |  |
| BA.4/BA.5 RBD | 4.35E+05 | 3.58E-03 | 8.22E-09 |  |  |  |  |  |  |
| XBB.1.1 RBD | *N.B.* | *N.B.* | *N.B.* |  |  |  |  |  |  |
| XBB.1.5 RBD | 4.63E+04 | 4.60E-03 | 9.93E-08 |  |  |  |  |  |  |
| BQ.1.1 RBD | *N.B.* | *N.B.* | *N.B.* |  |  |  |  |  |  |
| EG.5.1 RBD | 1.60E+03 | 2.89E-01 | 1.80E-04 |  |  |  |  |  |  |
| HV.1 RBD | 3.94E+01 | 3.70E-02 | 9.39E-04 |  |  |  |  |  |  |
| JN.1 RBD | 3.66E+01 | 8.44E-02 | 2.31E-03 |  |  |  |  |  |  |

**Supplemental Table 2.**
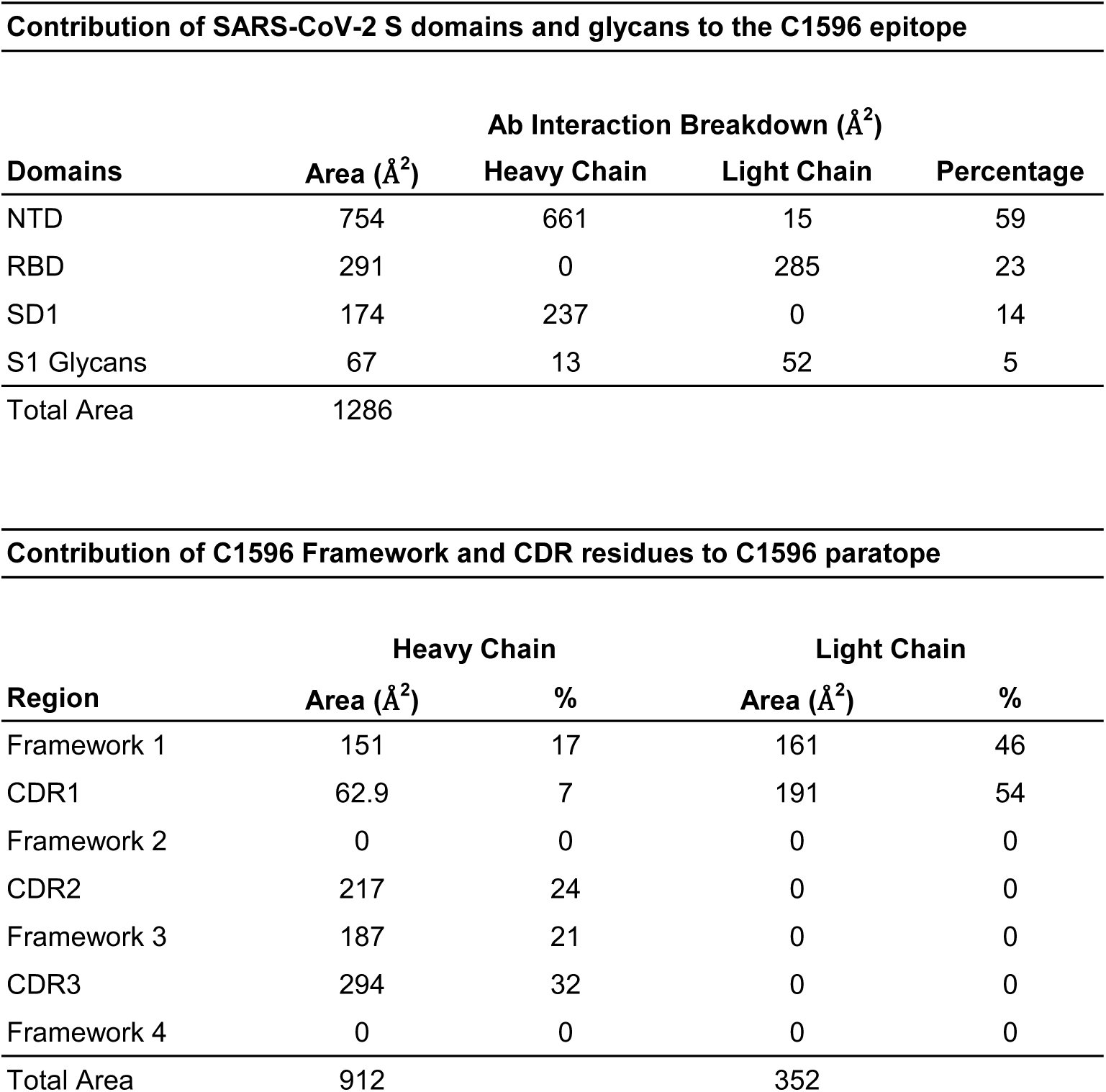
C1596 epitope and paratope buried surface area. Related to Figure 2.

**Contribution of SARS-CoV-2 S domains and glycans to the C1596 epitope**
| Domains | Area (Å<sup>2</sup>) | Ab Interaction Breakdown (Å<sup>2</sup>) |  |  |
| --- | --- | --- | --- | --- |
|  |  | Heavy Chain | Light Chain | Percentage |
| NTD | 754 | 661 | 15 | 59 |
| RBD | 291 | 0 | 285 | 23 |
| SD1 | 174 | 237 | 0 | 14 |
| S1 Glycans | 67 | 13 | 52 | 5 |
| Total Area | 1286 |  |  |  |

| Region | Heavy Chain |  | Light Chain |  |
| --- | --- | --- | --- | --- |
|  | Area (Å<sup>2</sup>) | % | Area (Å<sup>2</sup>) | % |
| Framework 1 | 151 | 17 | 161 | 46 |
| CDR1 | 62.9 | 7 | 191 | 54 |
| Framework 2 | 0 | 0 | 0 | 0 |
| CDR2 | 217 | 24 | 0 | 0 |
| Framework 3 | 187 | 21 | 0 | 0 |
| CDR3 | 294 | 32 | 0 | 0 |
| Framework 4 | 0 | 0 | 0 | 0 |
| Total Area | 912 |  | 352 |  |

**Supplemental Table 3.**
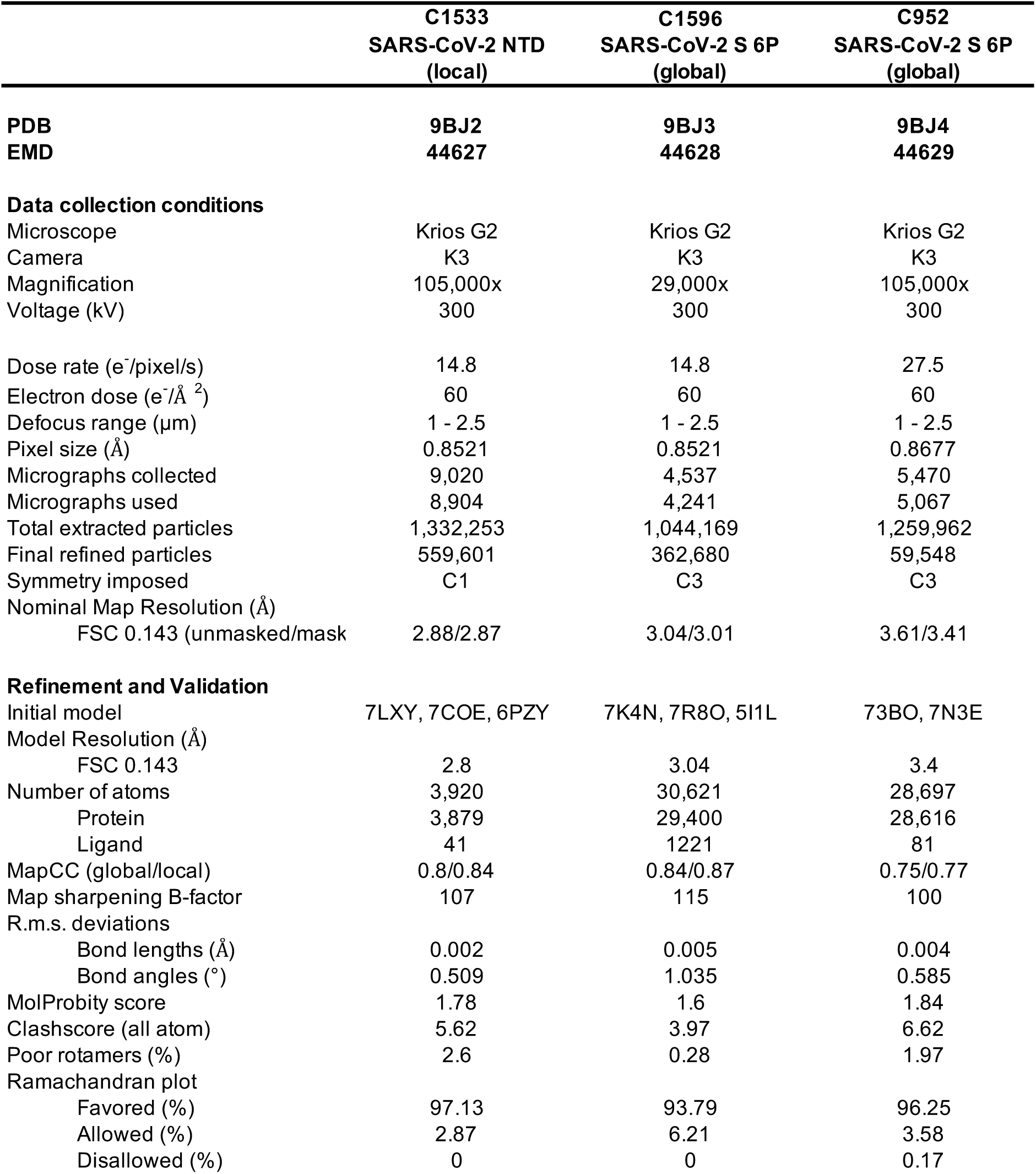
Cryo-EM data collection and data processing statistics. Related to Figures 2 and 3 and Supplemental Figure 1.

**Supplemental Table 4.**
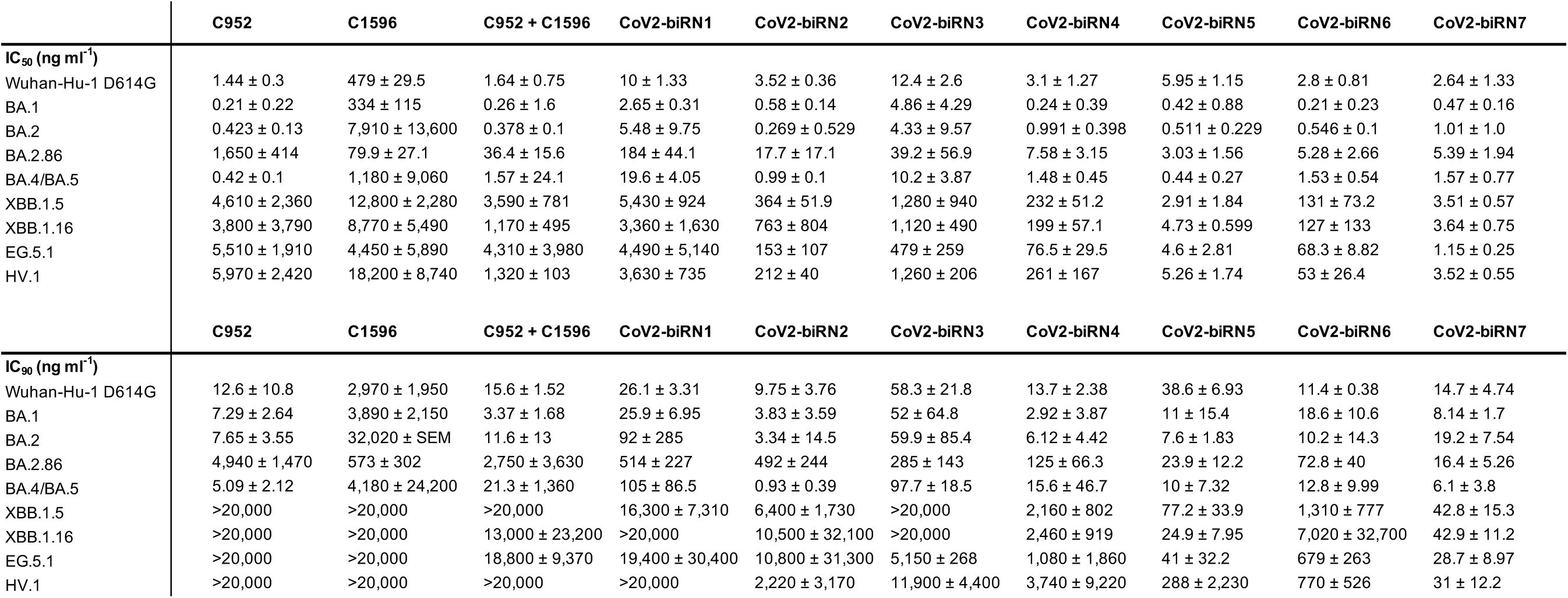
*In vitro* pseudovirus neutralization assay IC_50_ and IC_90_ values for monoclonal antibodies and CoV2-biRNs. Related to Figures 1 and 4.

## Notes

https://github.com/dms-vep/SARS-CoV-2_XBB.1.5_spike_DMS_Barnes_mAbs

